# Gelato streamlines reproducible and auditable assembly of western blot figures

**DOI:** 10.64898/2026.09.11.750878

**Authors:** Maria A. Pirozhkova, Elisheva Babitz, Daniel Benhalevy

## Abstract

**Summary:** Western blotting has been widely used for decades, yet figure preparation remains unstandardized, error-prone and difficult to audit. Gelato is a free Fiji/ImageJ plugin that streamlines intuitive figure preparation, logs processing coordinates, provides a straightforward audit platform and enables reproduction from raw images. By making these capabilities simple and accessible, Gelato facilitates robust Western blot reporting, reducing a substantial burden on authors and reviewers while strengthening scientific rigor.

**Availability and Implementation:** Gelato, source code and examples are freely available for download at https://github.com/masha-rgfj/Gelato-fiji-plugin. Implemented in Java and runs on any platform supported by Fiji. See README and manual for full documentation.

**Contact:** (M.A.P.), (D.B.)

**Supplementary information:** supplementary data is available at Bioinformatics online.

## 1 Introduction

Published science increasingly needs to be accompanied by sufficient source information for inspection and reproduction. This includes Western blot (WB) figures, although submission of source-images alone does not reveal the sequence of transformations linking the original data to published figures[1,2]. One of the main challenges is that the molecular weight (Mw) size markers and blot signals are recorded in separate image files, requiring the integration of information from the two acquisitions during cropping, rotation and resizing. One common approach is to overlay the colorimetric and chemiluminescent images (e.g. using ImageLab [Bio-Rad], QuickFigures[3] or Sciugo.com [Brainpoint Incorporated]). This preserves the exact positions of the Mw markers but introduces noise from the colorimetric image into the resulting composite, compromising appearance and quantification. An alternative common approach is to transfer Mw marker positions manually to the blot image using graphics software (e.g. Adobe CS, Inkscape or PowerPoint). This preserves a clean blot image but requires skill and careful safeguard of the correspondence between the two images through cropping, rotation, resizing, annotation, and panel assembly. These operations are thus labor-intensive and error-prone, and the relationship between the final figure, its source-image regions and the Mw coordinates used for labelling is not recorded, and difficult to inspect.

Auditing WB figures is evidently relevant, as some journals already require, in addition to uncropped blots, the submission of audit figures highlighting cropped regions of interest and Mw marker annotations[1]. In accordance, reproducibility concerns have given rise to dedicated WB analysis software, such as IOCBIO Gel[4], which safeguards densitometry of signal intensity. However, source-data submission and reproducible densitometry do not document figure assembly and size marker positioning. In addition, although essential, current practices for reviewing WB figures are suboptimal and inefficient, burdening authors, publishers and reviewers. Here, we present Gelato - A Fiji/ImageJ plugin that efficiently addresses Western blot reporting, from meeting authors need for precise and simple figure assembly, through background logging of transformations, to expediting figure inspection and reproduction.

## 2 Implementation and application

Gelato is a Fiji plugin written in Java. It requires the iText library for PDF generation, which is bundled with Fiji or can be added to ImageJ. Gelato can be installed by placing the .jar from the project’s GitHub repository into the Fiji Plugins folder, or by adding the Fiji update site (instructions in README).

Gelato guides the user through blot-figure assembly as an intuitive sequence of recorded operations (Fig 1). Its key features are matched marker-blot coordinate mapping, and logged figure production that can be reloaded for audit and reproduction.

**Figure 1.**
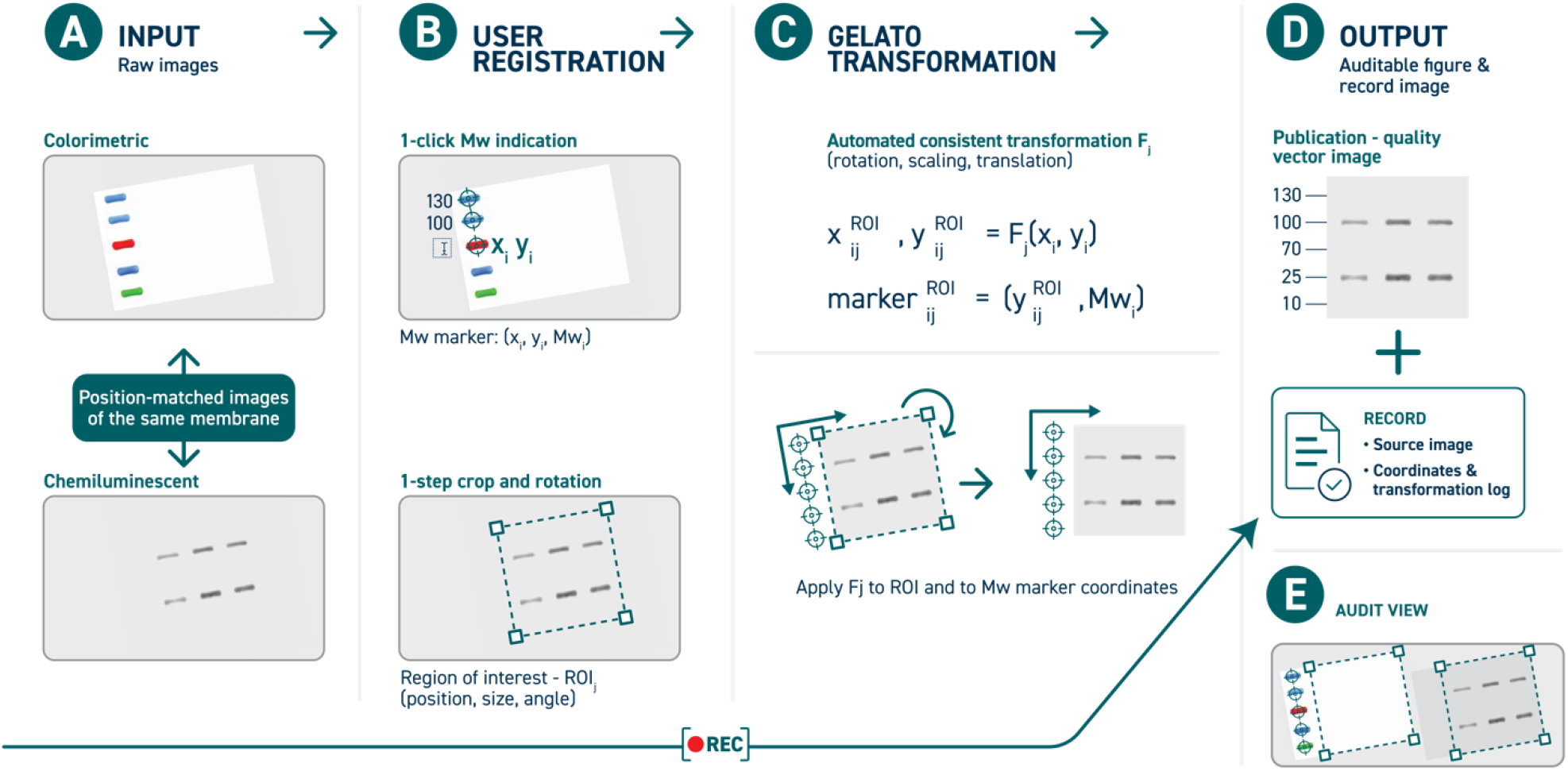
Gelato. **A. Inputs**. Colorimetric (top) and chemiluminescent (bottom) position-matched images of the same membrane/s. **B. User’s registration of size markers and regions of interest**. Each molecular weight (Mw) position is single-click marked, and size values are registered (top). Regions of interest (ROIs) crop and tilt in a single step per ROI (bottom). **C. Transformation**. Per ROI, relevant Mw data is automatically detected, and a to-horizontal rotation undergoes, keeping the correct geometry between the size marker and ROI. **D. Output**. A figure vector image is exported along with a log text file, enabling effective figure audit and reproduction from source images. **E. Audit view**. Upon submission of a log file to Gelato, raw image files are called by name, and then crops and Mw markers are highlighted on the raw images (see Supplementary Fig. 1 for authentic view including the reproduced final figure).

After user registration of molecular-weight (Mw) markers (Fig. 1B, top), and selection of region of interest (ROI) using a one-step crop and tilt (Fig. 1B, bottom), Gelato projects the Mw coordinates onto each selected ROI, accounting for tilt (Fig. 1C), and automatically attaches the corresponding Mw labels alongside the ROI (Fig. 1D). Mw marker labels are assigned by default to the left-hand side of each ROI, and an option for right side assignment can be selected. Since the marker and blot images have identical dimensions and share the same pixelwise-registered coordinate system, marker (i) is represented as:

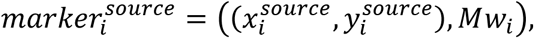

and ROI (j) is described by its origin, dimensions, and rotation angle:

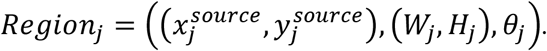

Coordinates follow the image convention in which (x) increases rightward, (y) increases downward, and *θ*_*j*_ is measured clockwise.

When the ROI crop is confirmed, it defines an affine transformation from source-image coordinates to the upright, ROI-local coordinate system:

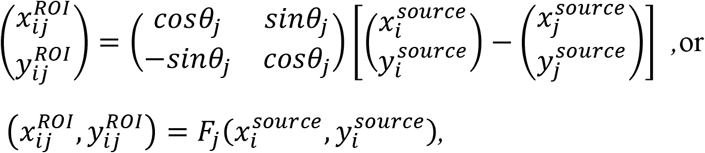

where *F*_*j*_ is the affine transformation defined in the equation above (Fig. 1C).

The raster ROI is rendered in this ROI-local coordinate system using bicubic interpolation, whereas marker coordinates are transformed analytically using the same geometric mapping. Consequently, the ROI-local vertical position of marker (i) is

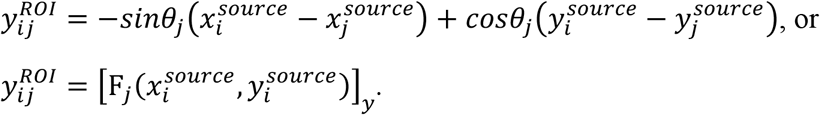

Here, []_*y*_ denotes the vertical component. Only the vertical component is required because each MW marker is horizontally translated to the ROI side. If the ROI is later resized, the marker’s position on canvas is scaled consistently. Their geometric registration is therefore maintained after ROI deskewing and panel resizing, eliminating the need for manual marker repositioning (Fig 1C-D).

Coupled to figure production, Gelato exports a readable coordinates log that contains all the information required for figure audit and reproduction from co-submitted raw images (Fig 1D, and Supplementary information - Manual). For each ROI, the log states the source image, its dimensions, the crop coordinates, used marker sets and their coordinates.

In reconstruction (audit) mode, relationship between declared source images and the assembled figure is made directly inspectable (Fig 1E). First, user loads the log file, and Gelato requests every image stated in the log. Each crop is regenerated, Mw marker positions are recalculated from the referenced marker set and projected into the rotated crops, and Gelato displays ROIs crop boundaries and original Mw marking on the raw images, as well as a reconstructed final figure. This reconstruction intentionally omits text labels. If needed, annotated views can be exported as full-resolution PNG images or PDFs containing the source raster with vector overlays.

Gelato performs consistency checks on the supplied source images and reconstruction record. Mismatches in image dimensions and aspect ratios are shown as warnings that require confirmation before proceeding, leaving assessment with the user or reviewer. Source-image paths are recorded for convenience, but reconstruction is based on Gelato requesting the user to supply raw images (by file name), allowing records and source files to be moved or transferred.

For additional robustness, the reconstruction tolerates partial and externally generated records. Crop geometry can be represented either by three reference points or by an origin together with crop dimensions and rotation angle. Gelato records both representations by default, allowing either representation to be used independently for reconstruction. If both are present, discrepancies exceeding one source pixel require the user to select which geometry to use. This redundancy enables reconstruction from records generated outside Gelato, provided that one complete geometric representation is supplied. Similarly, when the subset of molecular-weight markers used for a crop is not specified explicitly, Gelato reconstructs all registered markers whose mapped positions fall within the crop range. All inferred information, used geometry and warnings are summarized in a reconstruction report. This way Gelato reproduces the declared assembly steps so that the correspondence can be evaluated directly by the reviewer with or without algorithmic judgement.

## 3 Discussion

The value of maintaining an explicit connection between published western blot figures and their source images extends beyond convenience. A systematic assessment of 551 publications from top-quartile journals found that more than 90% presented only cropped blots, more than 80% did not provide raw images, and more than 95% did not indicate molecular-weight markers[1]. Image manipulation was the most common issue identified in a systematic analysis of PubPeer comments[5], while journal-level screening has reported missing corresponding raw blots, undisclosed image reuse and different crops of the same image presented as distinct experimental conditions[6]. Importantly, such observations do not necessarily indicate misconduct, because inappropriate image duplication can also arise from unintentional errors during figure preparation[7]. In either case, they illustrate the value of preserving the connection between source images and the final figure during figure preparation, rather than attempting to reconstruct it retrospectively.

This concern is not new. The need to preserve the relationship between source data and figures was recognized more than a decade ago, with FigureJ introduced in 2013 as a structured approach to reproducible multi-panel figure construction[8]. More recently, reproducibility concerns have motivated dedicated analysis software that records source images together with analytical operations, allowing quantitative analysis to be reproduced[4,9,10]. Gelato complements these efforts by applying the same principle of simultaneous recording to the final figure assembly stage. Existing tools support important parts of the processing, including figure layout, annotation, densitometry and automated line detection (Supplementary Table 1), but the WB specific combination of marker and blot matched images, explicit Mw-coordinate correspondence and a possibility to review the figure assembly remains insufficiently addressed.

Importantly, Gelato-generated output is also readily available for computational inspection by other tools, including AI agents. However, for experimentalists, AI-based agents cannot currently replace personal liability for proper image processing. Thus, as in other cases requiring human accountability, Gelato’s approach provides a practical, effective and future-proof platform for WB reporting. By coupling simple WB figure preparation with automatic recording of source assignments, coordinates and transformations, Gelato turns an otherwise largely undocumented sequence of image-processing decisions into a streamlined inspectable workflow, by both human and machines.

## Supporting information

Gelato 1.1 User Manual

## Author contributions

Maria Pirozhkova (Conceptualization, Software, Writing – original draft; review and editing), Elisheva Babitz (Software, Validation,), Daniel Benhalevy (Supervision, Conceptualization, Writing – review and editing)

## Supplementary Material

Supplementary Figure 1 (Gelato audit view), Gelato manual, and Supplementary Table 1 (comparison with existing software) are provided online.

## Declaration of Generative AI usage

Open AI Codex (the GPT -5.6 Sol model) was used to develop and extend the Java implementation based on the author’s original Python standalone script. The authors directed the software design and functionality, and reviewed, tested, debugged, and revised the generated code. The authors take full responsibility for the final implementation and its validation. Additionally, the model was used to refine the text of the manuscript.

## Data availability

Gelato source code, along with example raw images, output figures and logs, is publicly available on GitHub (https://github.com/masha-rgfj/Gelato-fiji-plugin) and archived on Zenodo (10.5281/zenodo.23766971). Gelato is distributed under the GNU General Public License v3.0.

## References

1. Kroon C, Breuer L, Jones L, et al. Blind spots on western blots: Assessment of common problems in western blot figures and methods reporting with recommendations to improve them. PLOS Biology. 2022;20(9):e3001783. doi:10.1371/journal.pbio.3001783

2. Janes KA. An analysis of critical factors for quantitative immunoblotting. Science Signaling. 2015;8(371):rs2–rs2. doi:10.1126/scisignal.2005966

3. Mazo G. QuickFigures: A toolkit and ImageJ PlugIn to quickly transform microscope images into scientific figures. PLOS ONE. 2021;16(11):e0240280. doi:10.1371/journal.pone.0240280

4. Kütt J, Margus G, Kask L, et al. Simple analysis of gel images with IOCBIO Gel. BMC Biology. 2023/10/20 2023;21(1):225. doi:10.1186/s12915-023-01734-8

5. Ortega JL. Classification and analysis of PubPeer comments: How a web journal club is used. Journal of the Association for Information Science and Technology. 2022;73(5):655–670. doi:10.1002/asi.24568

6. Williams CL, Casadevall A, Jackson S. Figure errors, sloppy science, and fraud: keeping eyes on your data. The Journal of Clinical Investigation. 03/25/ 2019;129(5):1805–1807. doi:10.1172/JCI128380

7. Bik EM, Fang FC, Kullas AL, Davis RJ, Casadevall A. Analysis and Correction of Inappropriate Image Duplication: the Molecular and Cellular Biology Experience. Molecular and Cellular Biology. 2018/10/01 2018;38(20):e00309–18. doi:10.1128/MCB.00309-18

8. Mutterer J, Zinck E. Quick-and-clean article figures with FigureJ. J Microsc. Oct 2013;252(1):89–91. doi:10.1111/jmi.12069

9. Gulbulak U, Wellette-Hunsucker AG, Kampourakis T, Campbell KS. GelBox: open-source software to improve rigor and reproducibility when analyzing gels and immunoblots. American Journal of Physiology-Heart and Circulatory Physiology. 2024;327(3):H715–H721. doi:10.1152/ajpheart.00144.2024

10. Sanders MT. BlotTool: A reproducible command-line workflow for western blot densitometry. SoftwareX. 2026/09/01/ 2026;35:102859. doi:10.1016/j.softx.2026.102859

