## Supplementary material for "Gelato streamlines reproducible and auditable assembly of western blot figures": Gelato 1.1 User Manual

### About

Gelato is a Fiji/ImageJ plugin for assembling gel figures while keeping a clear, human-readable record of every source image, crop, and molecular-weight marker that can be used for instant audit.

The source code, the plugin, and example files are available at https://github.com/masha-rgfj/Gelato-fiji-plugin

Contact:

Gelato version 1.1.5 is archived on Zenodo: https://doi.org/10.5281/zenodo.23766971

### Installation

Gelato is designed for Fiji. It also runs in ImageJ when the iText module is available.

1. Download gelato-1.1.x.jar from https://github.com/masha-rgfj/Gelato-fiji-plugin latest release
2. Place gelato-1.1.x.jar in the Fiji plugins folder (Fiji.app/plugins).
    a. On macOS, open the Fiji application contents first, then place the file in the plugins directory.
3. Open Fiji.
4. Click Plugins on application menu and choose Gelato.

### Getting started

Supported image formats: TIFF, PNG, and JPEG. Gelato converts loaded images to RGB for consistent display and export.

You can build a new figure or reconstruct a figure from a coordinate log and source images.

### Figure creation

The plugin initially shows an A4-sized canvas where the figure is constructed, with a menu on the left side. Loaded images are opened in separate windows on the right side.

The workflow consists of three steps per images set: kDa marker registration, crop region selection, and annotation. After this streamlined process, ready figure can be exported in pdf, in which all objects beside the original image remain vector.

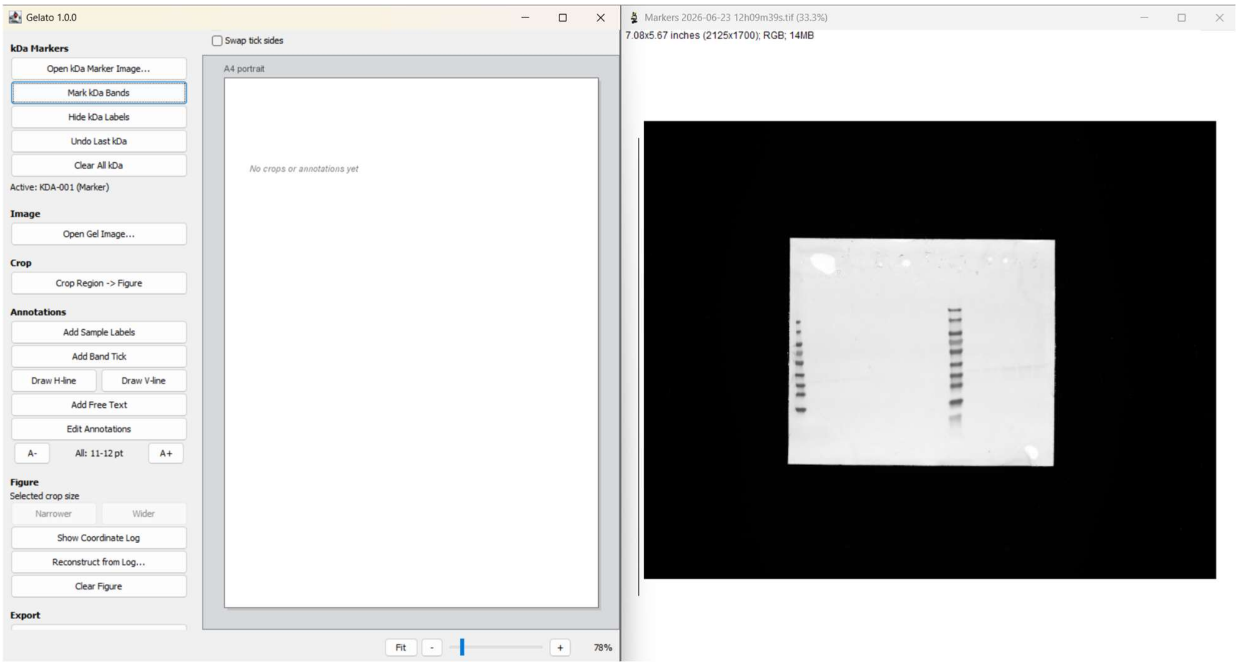

### kDa markers registration and crop region selection

Gelato works with paired marker-blot images without needing to directly overlay them, and with images where the marker and blot were pre-overlayed.

#### Workflow for Separate marker and blot images

1. kDa marker registration
   a. Click "Open kDa Marker Image…". Select the image. The image will appear on the right.

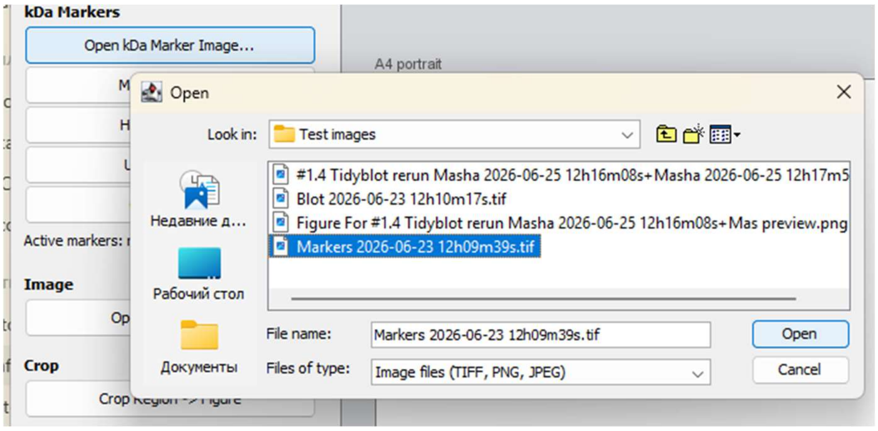

   b. On the marker image, click on a marker band and enter the Mw value. Continue until all desired marker bands are registered, for the entire set of vertically positioned ROIs ("see workflow example – multi-membranes images" below for horizontally positioned ROIs). "Undo Last kDa" or "Clear All kDa" on the left side menu can be used.

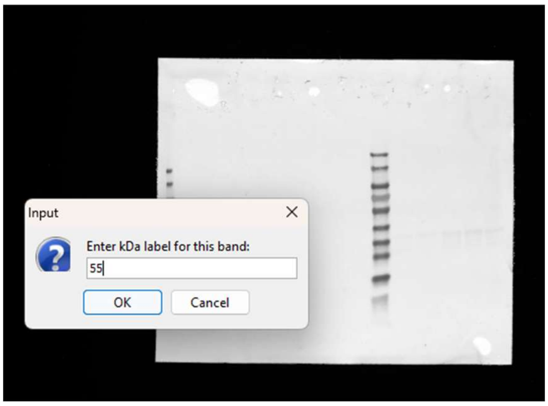

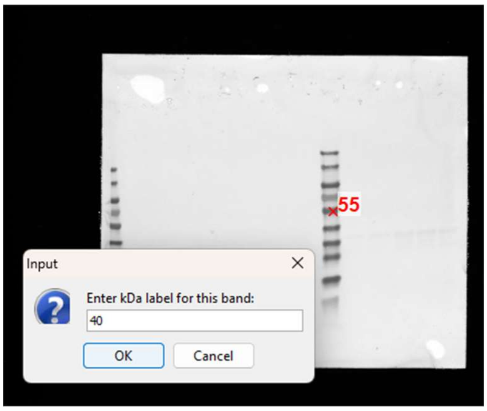

2. Crop regions of interest (ROIs)
    a. Click "Open Gel Image…". The marked positions will be indicated on the loaded image. "Show/Hide kDa bands" button controls kDa values indication.

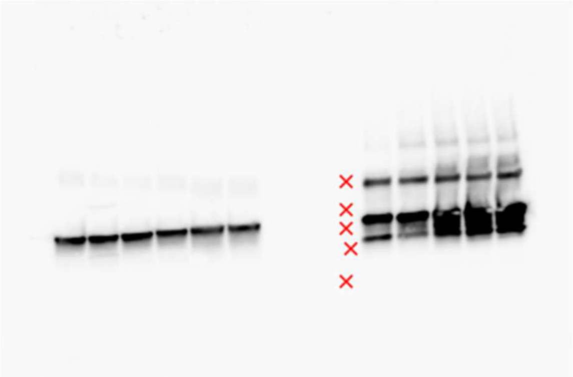

    b. Click *"Crop -> "* and select the region of interest by dragging and tilting the interactive rectangle.

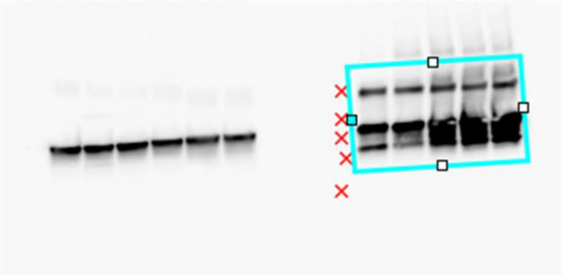

c. Click “Confirm Crop” and enter the label for the crop (e. g., protein name).

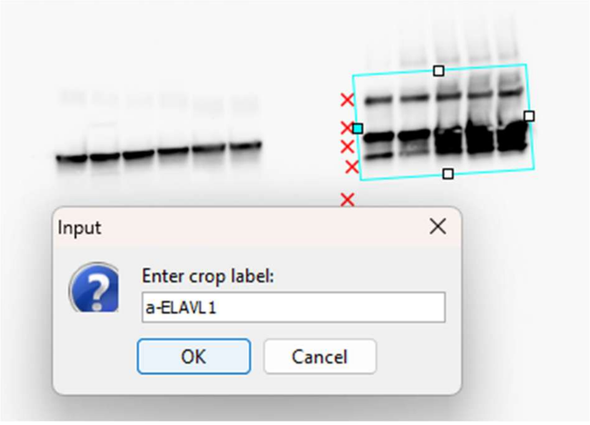

d. Click OK. The crop will appear on the A4 canvas together with registered kDa labels.

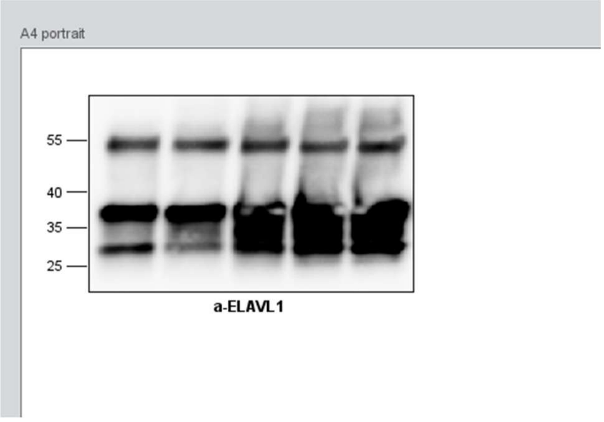

e. Repeat b-d for the full set of vertically positioned ROIs on the image.

If a new blot image is loaded without loading a new matched marker image, the same markers from step 1 will be applied to it (relevant for expedited preparation of the same figure from different exposure times of the same membrane). For a new images set, you can register a new marker set by repeating step 1 from opening a kDa Marker image. As long as Gelato session remains, transformations are recorded to a single log file for multiple images sets and ROIs, and all generated figure panels appear, vertically aligned, on the same canvas (panels can be resized with Narrower/Wider buttons, or moved and reordered by simple dragging).

#### Overlayed marker-blot image

Steps 1 and 2 slightly differ from the separate image case above: skip “Open kDa Marker Image…” and mark kDa bands on the loaded Gel Image directly.

1. kDa marker registration
   a. Click “Open Gel Image…”. Select the image. The image will appear on the right.
   b. Click “Mark kDa Bands”. On the Gel Image, click on a marker band and enter the Mw value. Continue until all desired marker bands are registered. “Undo Last kDa” or “Clear All kDa” can be used.

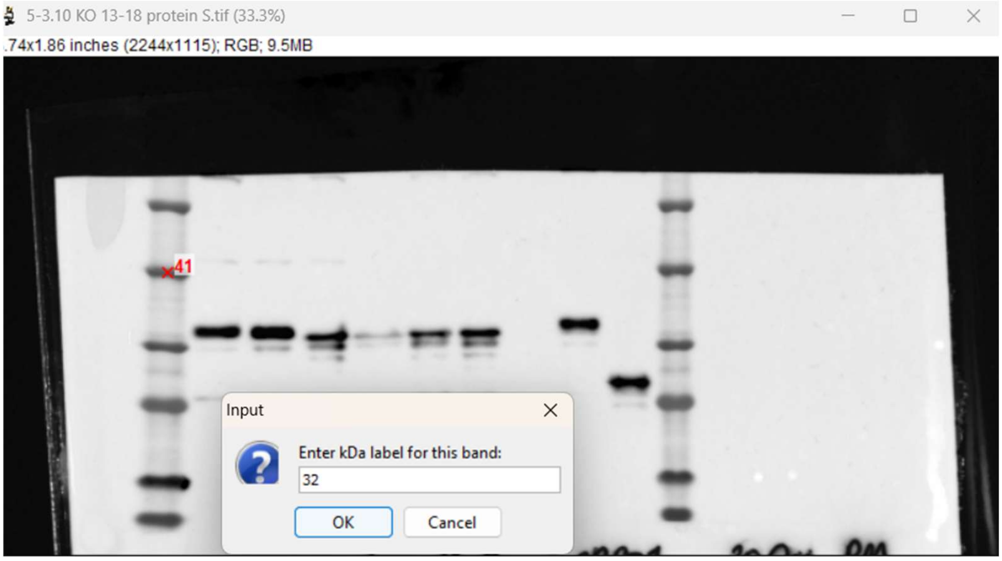

2. Crop ROIs
   a. Select each ROI by dragging and rotating the interactive rectangle. “Show/Hide kDa bands” button controls kDa values indication.

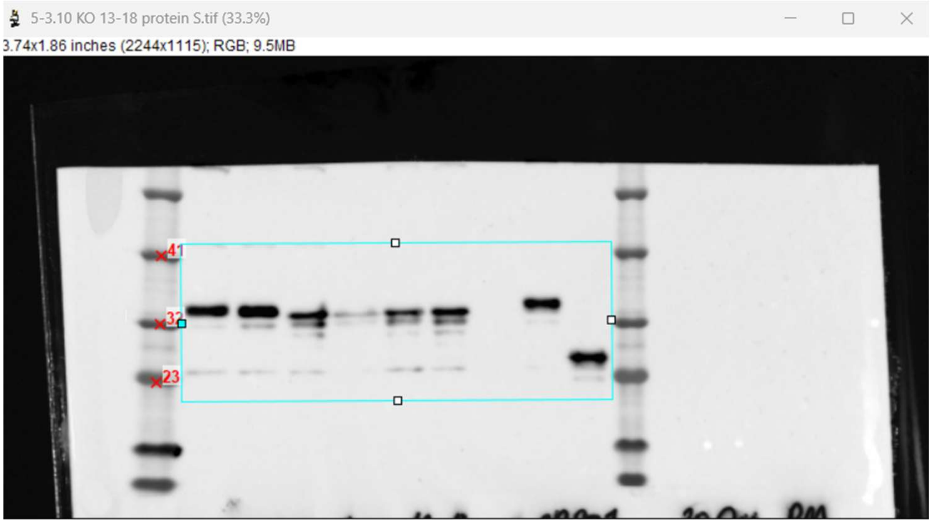

   b. Click “Crop Region -> Figure” and enter the label for the crop (e. g., protein or antibody name). The crop will appear on the A4 canvas together with registered kDa labels.

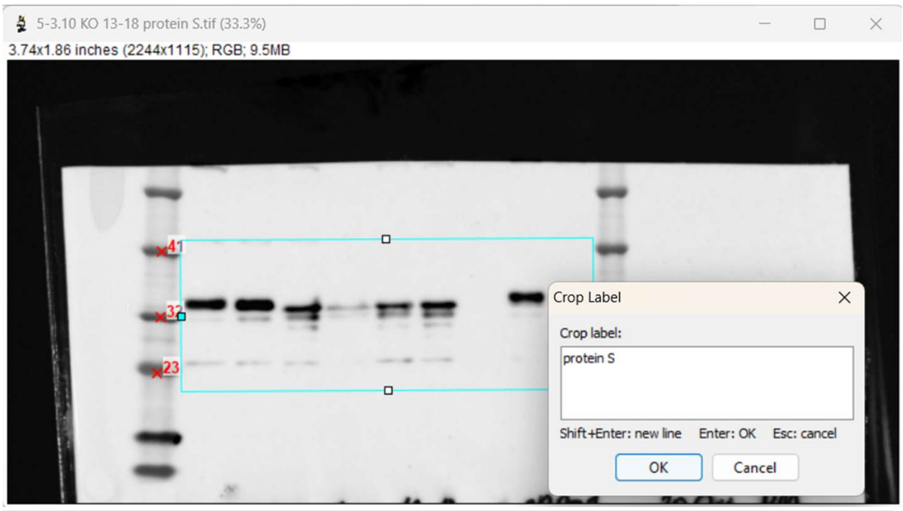

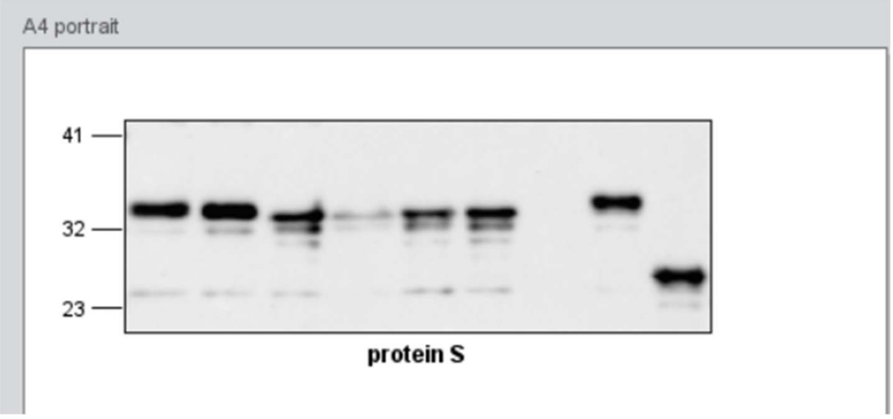

Repeat a-b for all ROIS on the image. If a new image is loaded, the same markers will be applied to it (expediting assembly of the same figure from different exposure times). You can register a new marker set by repeating step 1. The figure panel will appear one after the other on the canvas, vertically aligned, and can be moved and reordered by dragging.

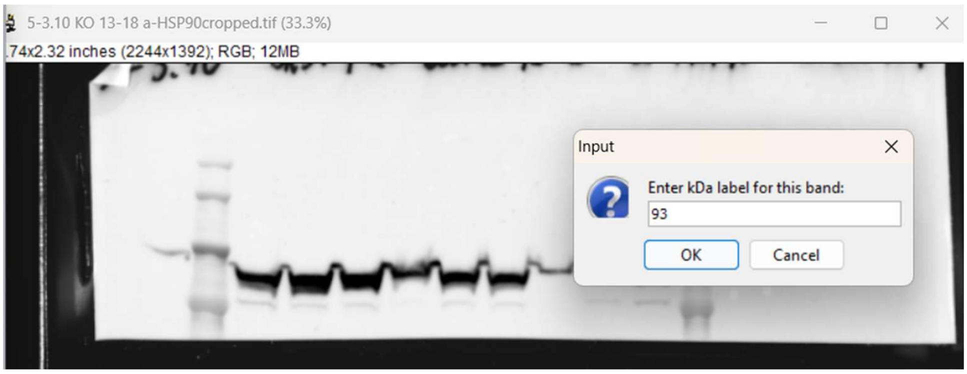

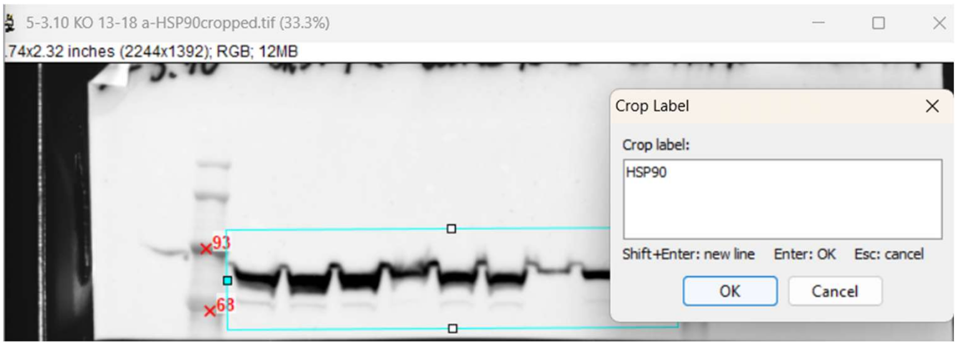

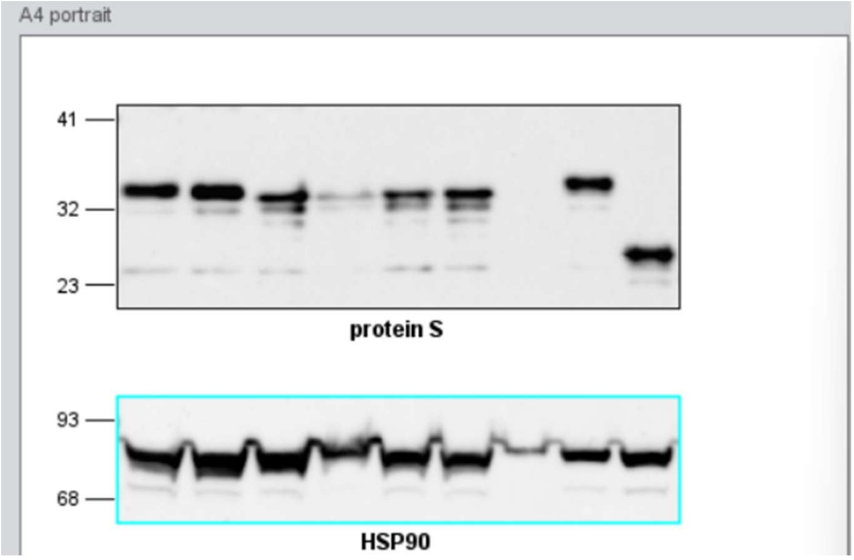

#### Workflow example - multi-membrane images

A single marker image can contain several independent marker sets. Each marker set represents one vertical series of Mw reference positions that can be linked to one or more ROIs.

Markers within the same set may extend vertically across the image. Therefore, if several membranes are positioned above or below one another, all their markers can be registered in a single step that will be linked to all the corresponding selected ROIs. In other words, all vertically spread marker sets are treated as a single marker set. However, as often relevant when multi-membranes are imaged, horizontally separated marker series will need to be indicated sequentially, though during the same Gelato session, and reporting to the same log file.

This example shows how to create a figure from matched marker and blot images containing six membranes that are spread both vertically and horizontally.

1. Registration of the first kDa marker set (three membranes on the left)
   a. Click “Open kDa Marker Image…”. Select the image. The image will appear on the right.
   b. Register the marker bands belonging to all vertically spread marker series. Click on a marker band and enter the Mw value. Continue until all desired marker bands are registered. “Undo Last kDa” or “Clear All kDa” on the left side menu can be used.

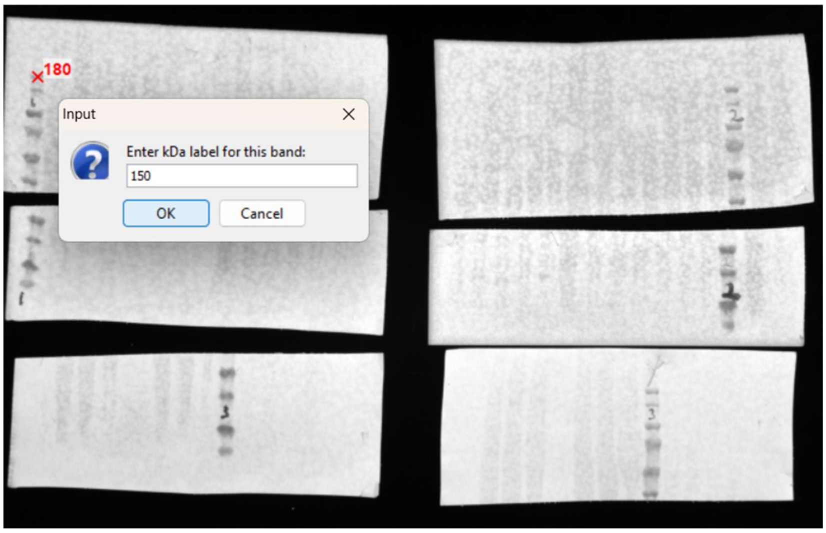
180
Input
Enter kDa label for this band:
150
OK
Cancel

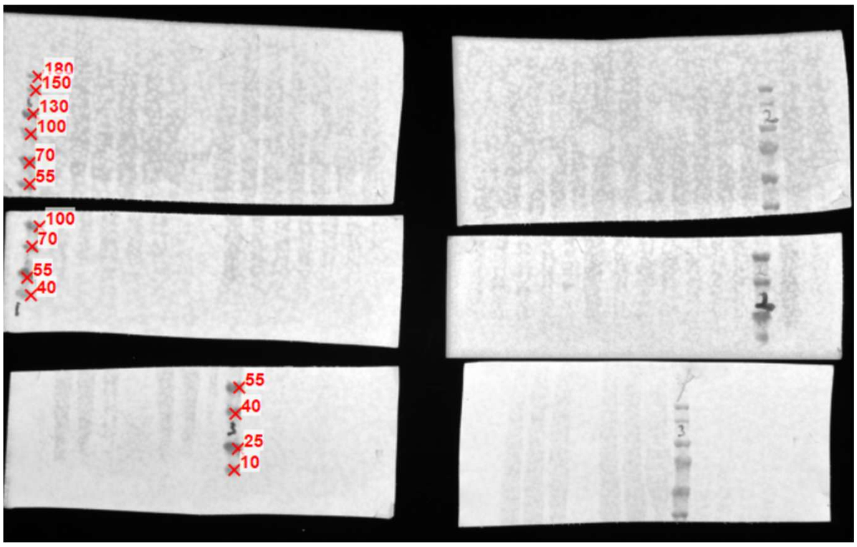
180
150
130
100
70
55
100
70
55
40
55
40
25
10

2. Selection of ROIs corresponding to the first kDa markers set

a. Click “Open Gel Image…”. The marked positions will be indicated on the loaded image. “Show/Hide kDa bands” button controls kDa values indication.

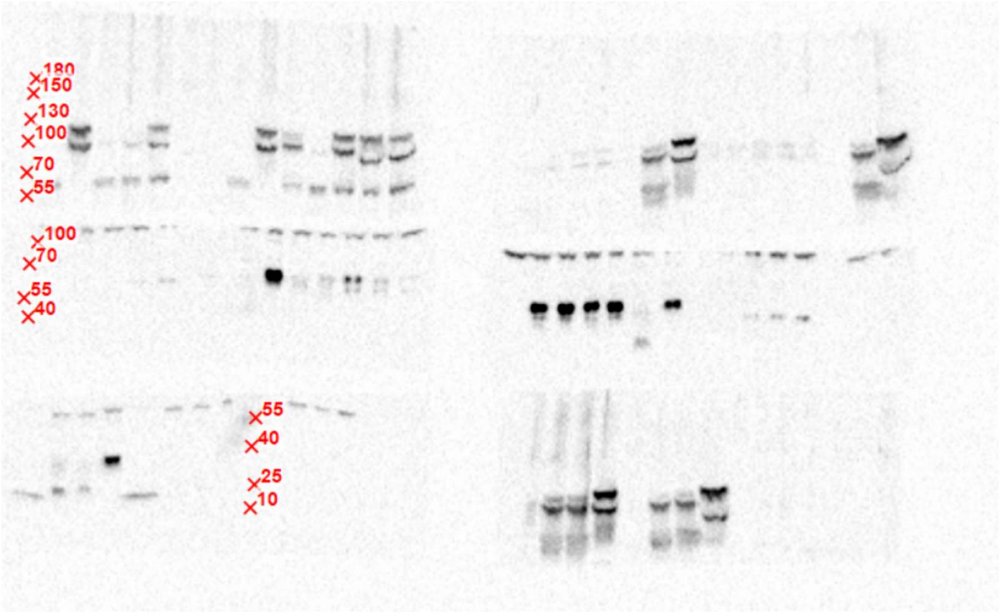

b. Select the region of interest by dragging the interactive rectangle. Click “Confirm Crop” and enter the label for the crop. The crop will appear on the A4 canvas together with registered kDa labels.

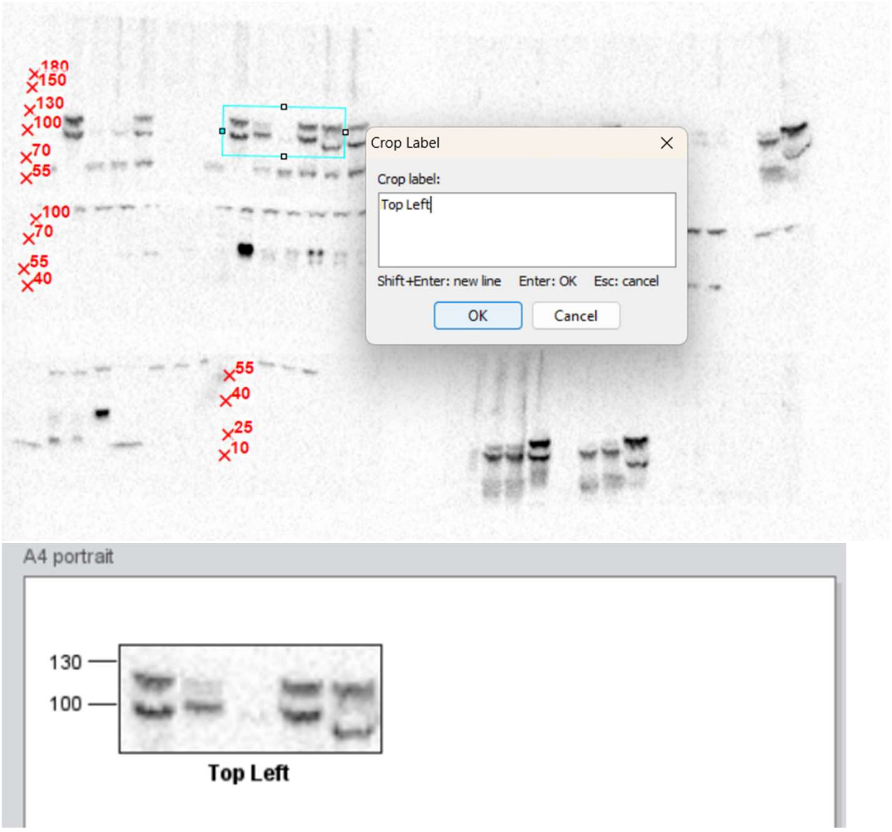

c. Repeat step b for all ROIs to which the indicated markers should be applied. The crops will appear one after another on A4 canvas, vertically aligned.

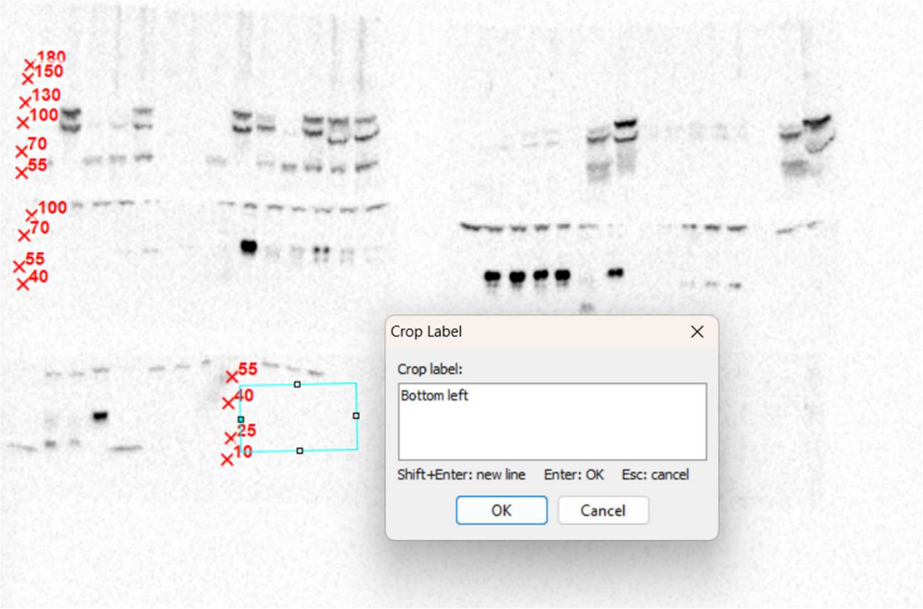

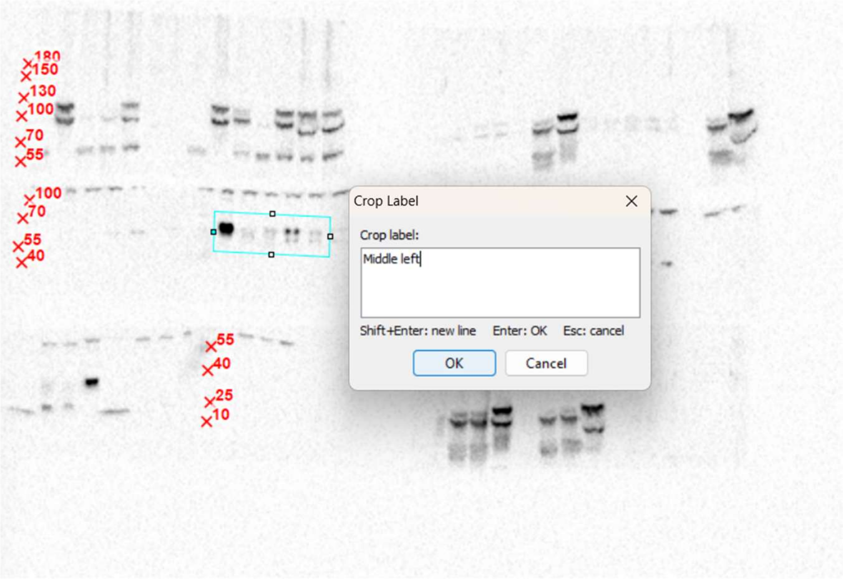

d. The crops can be moved and reordered by dragging and resized with Narrower/Wider buttons.

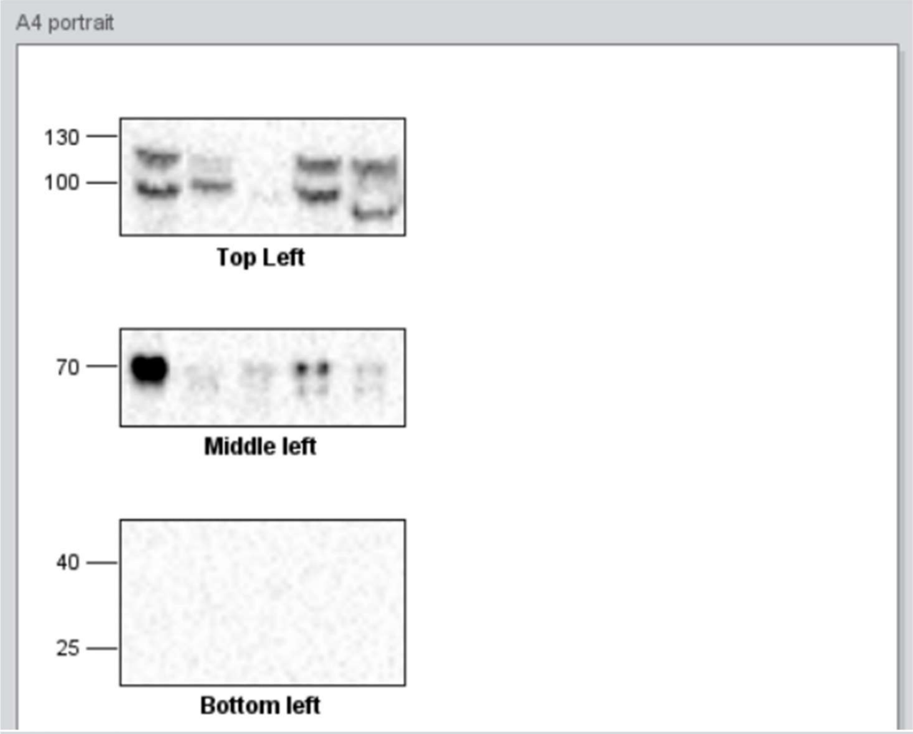

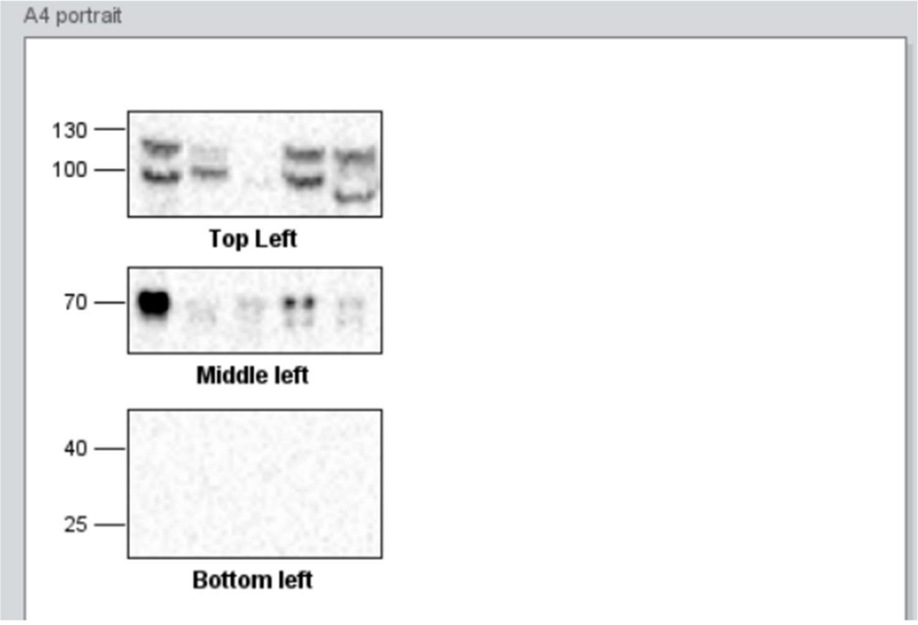

2. Registration of the second marker set
    a. When all ROIs that use the first indicated marker sets were processed, click “Open kDa Marker Image…” and select the same marker image again.
    b. Register the marker bands belonging to the second vertical marker series. Click on a marker band and enter the Mw value. Continue until all desired marker bands are registered. “Undo Last kDa” or “Clear All kDa” on the left side menu can be used.

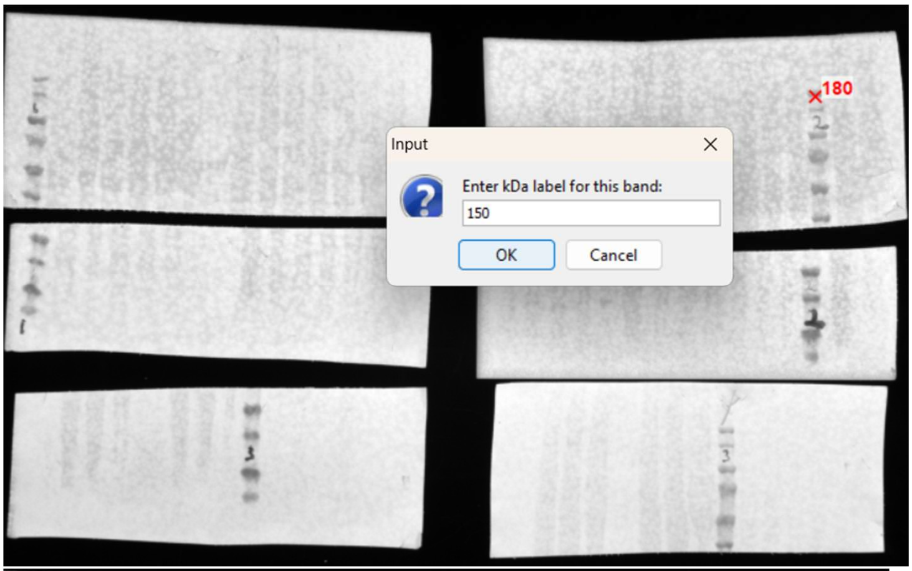

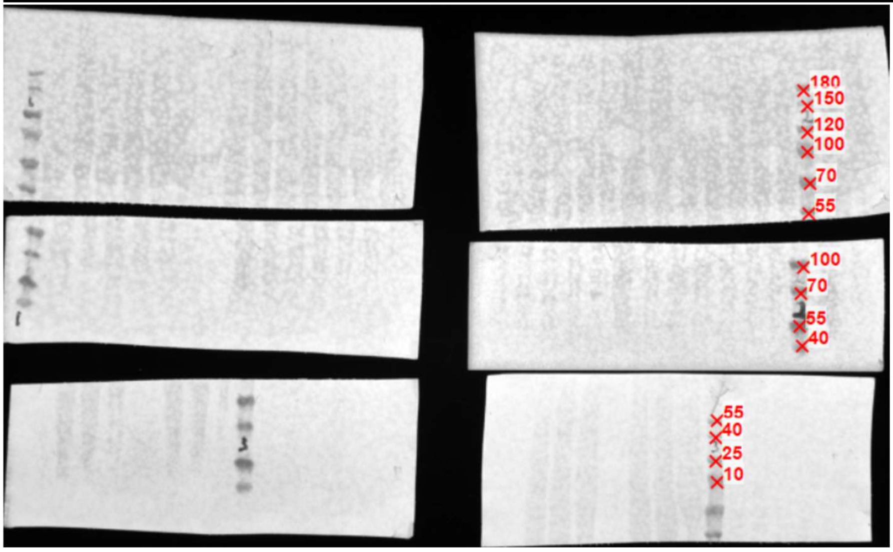

3. Selection of ROIs corresponding to the second kDa marker set
    a. Click “Open Gel Image…”. The positions from the last kDa marker set will be indicated on the loaded image. “Show/Hide kDa bands” button controls kDa values indication.

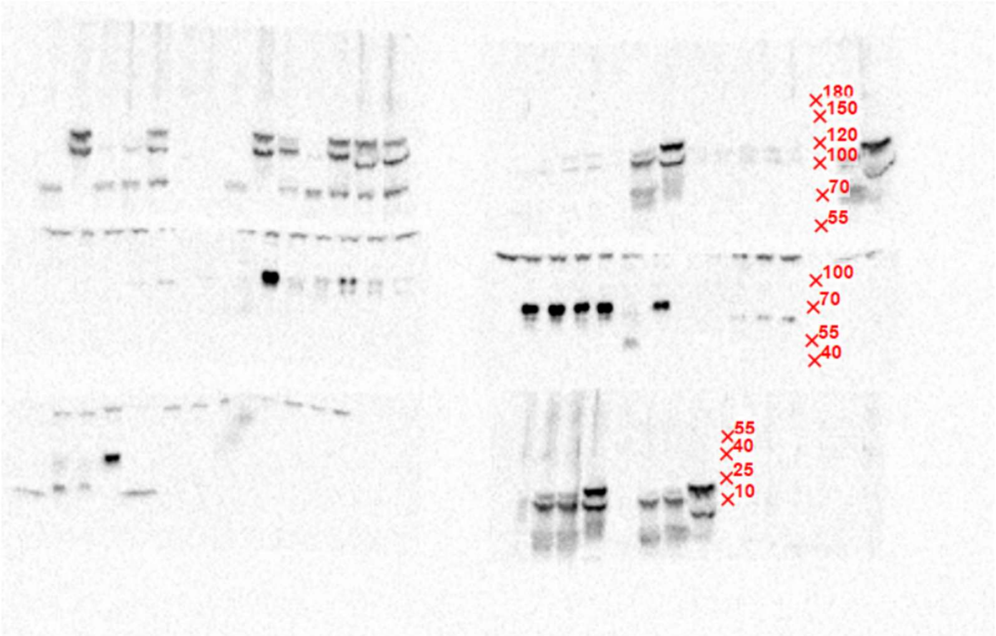

b. Select the region of interest by dragging the interactive rectangle. Click “Confirm Crop” and enter the label for the crop. The crop will appear on the A4 canvas together with registered kDa labels.

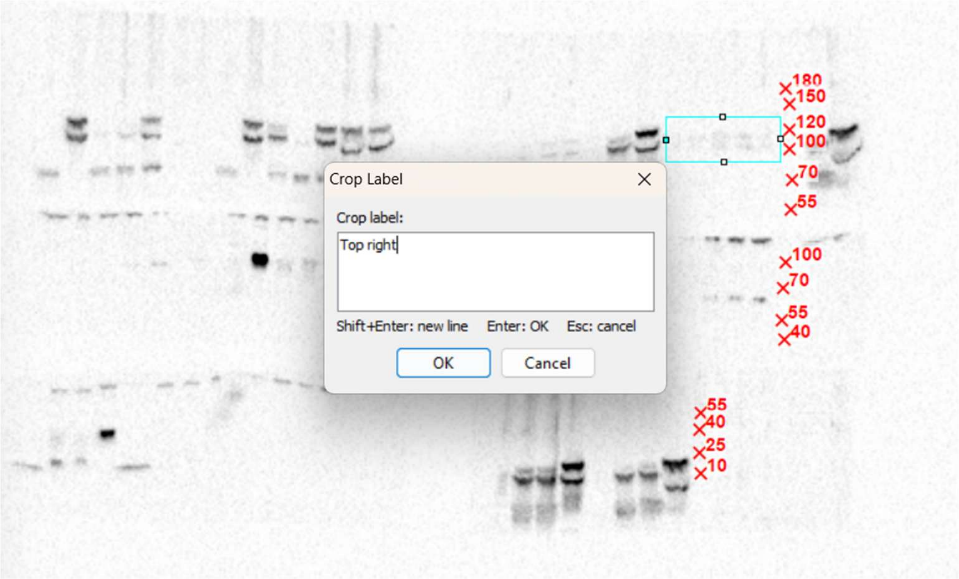

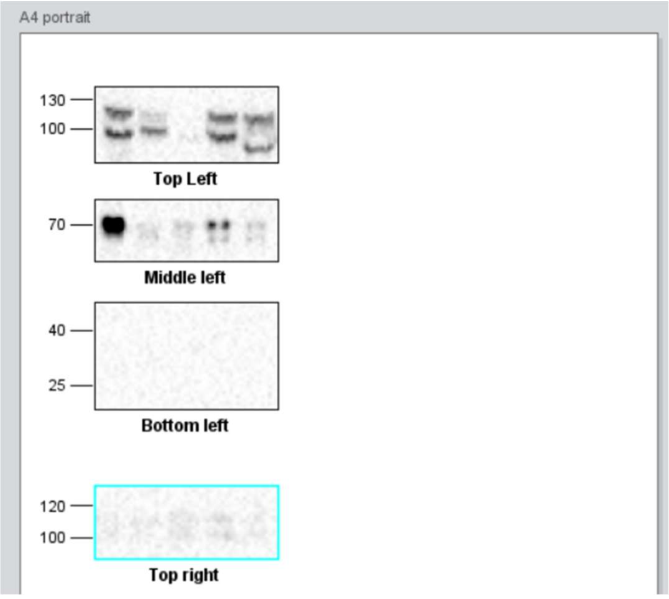

c. Repeat step b for all ROIs to which the indicated markers should be applied. The crops will be vertically aligned on A4 canvas.

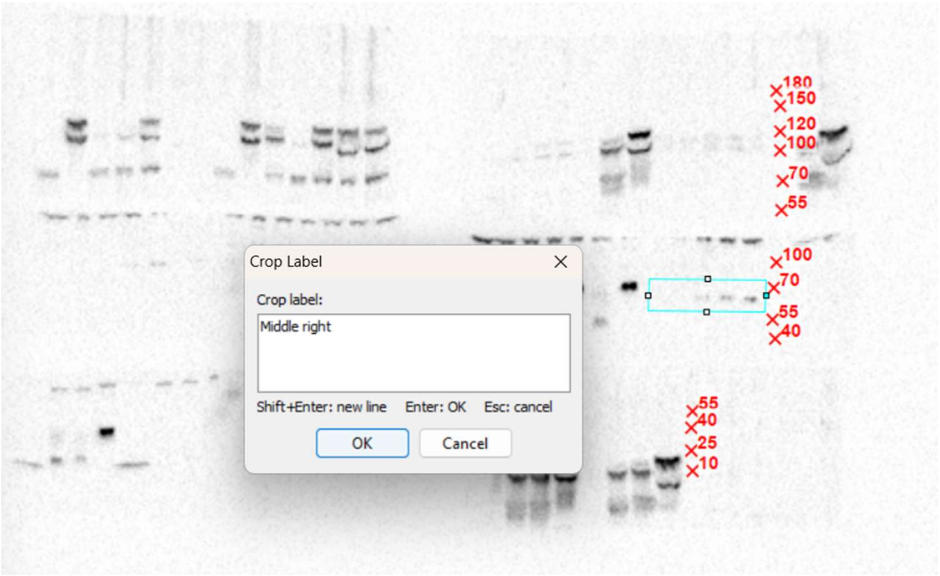

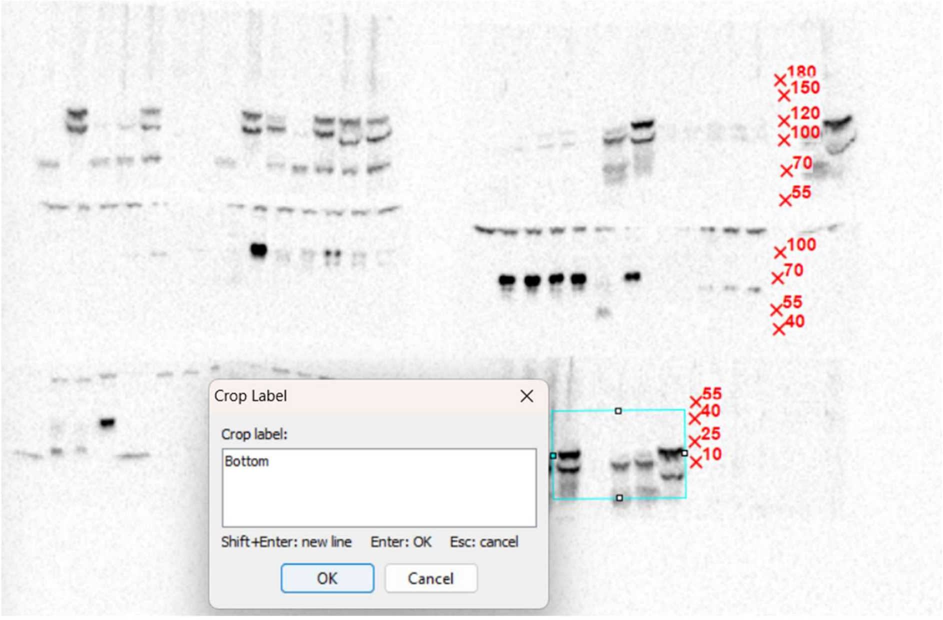

d. Crops can be rearranged by dragging or resized with Narrower/Wider buttons.

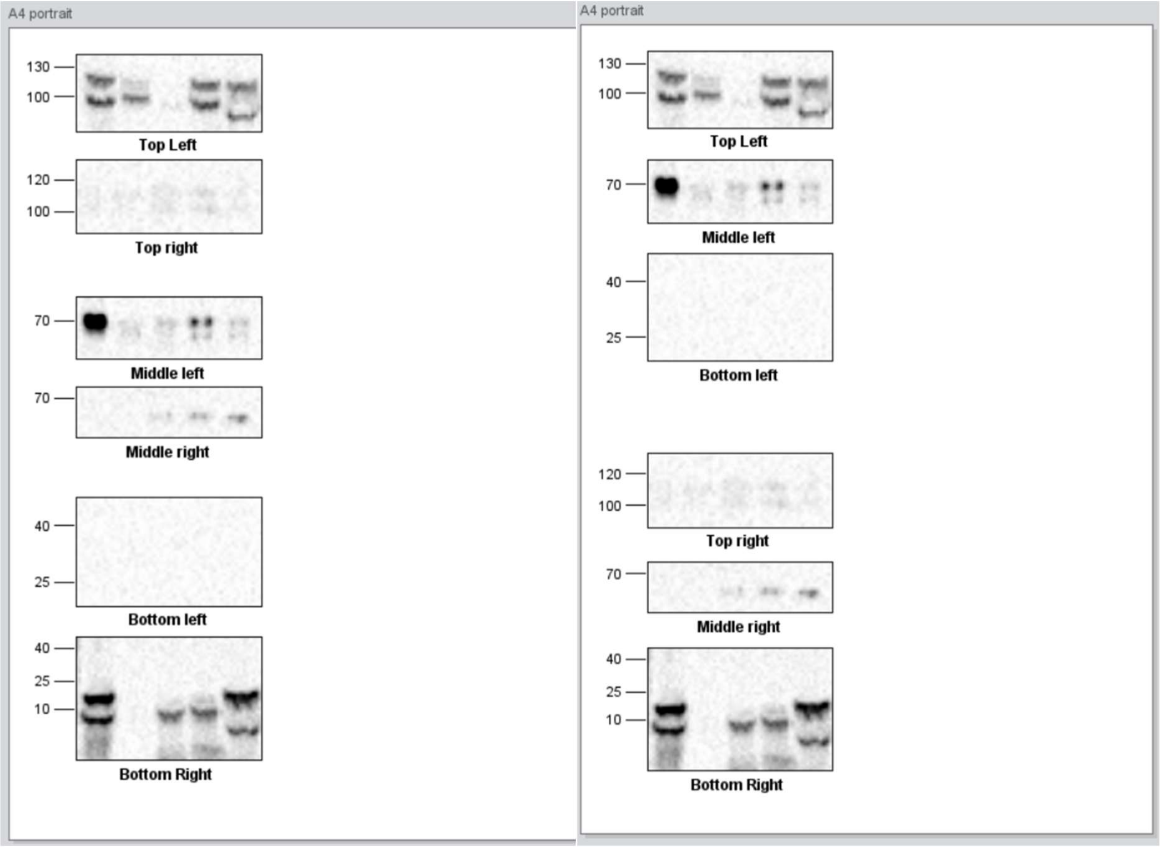

### Add annotations

Once ROIs and corresponding size markers are arranged on the canvas, additional annotations are enabled by Gelato. All objects remain independent vector for further processing. On Gelato, Molecular weight markers, Crop names, Sample labels and Band ticks are moved and scaled with the ROIs with font size preservation.

a. Click “Add Sample Labels”. Click on a lane within the ROI and enter Sample name. The name will appear above the selected point on top of the ROI. Continue until all lanes are labelled and then click “Stop Adding Labels”.

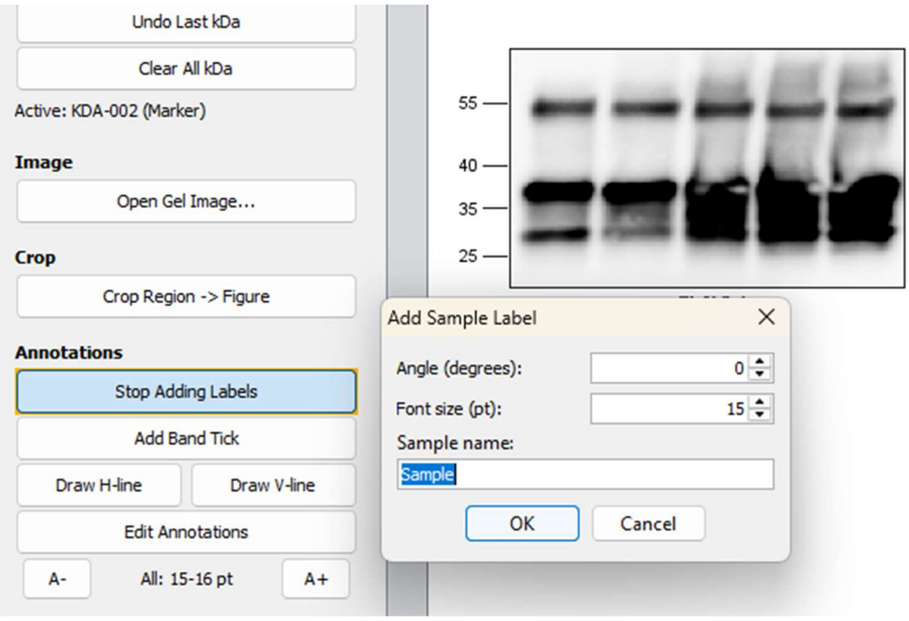

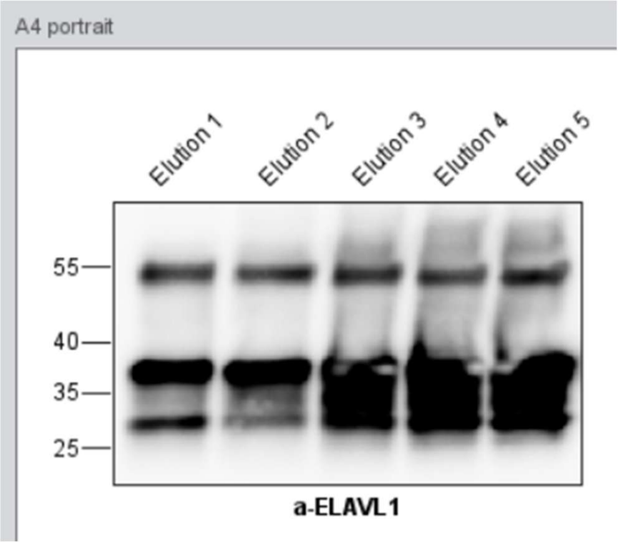

b. To highlight a particular band on the crop, click “Add Band Tick”

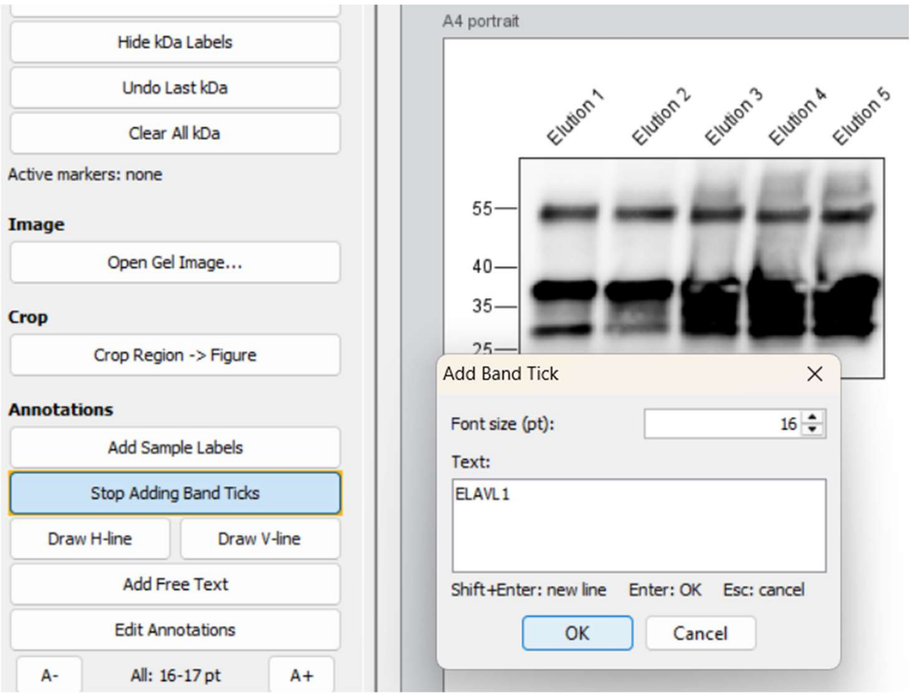

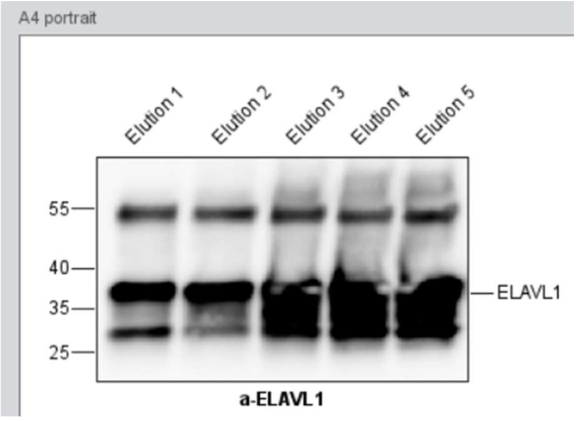

c. To add horizontal/vertical line, click “Draw H-/V- line”. Continue adding lines or click “Stop Drawing…”.

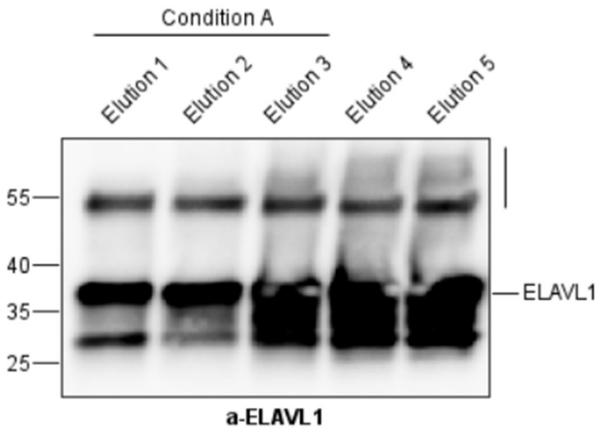

### Edit Annotations

After annotations are roughly inserted, they can be further refined, while the Mw markers are safely anchored to their positions.

Click “Edit Annotation” to start the Edit Annotations mode. When this mode is on, click on any annotation to select it (multi-selection is possible with Shift key). Possible changes will be listed on left-bottom edge of the window (e.g, “MW values are resize-only, use “A-/A+”). If nothing is selected, all text can be resized simultaneously by A+/A- buttons.

| | Mw markers | Crop names | Sample labels | Band ticks | H-/V- lines |
|---|---|---|---|---|---|
| Resize | A-/A+ | | | | Drag ends |
| Move (Drag/arrows) | No | Free | | Vertical | Free |
| Align/Distribute (right click) | No | Priority to Sample labels | | No | No |
| Text | No | Double-click | | | - |
| Angle | - | - | Double-click | - | - |

Editing horizontal and vertical lines

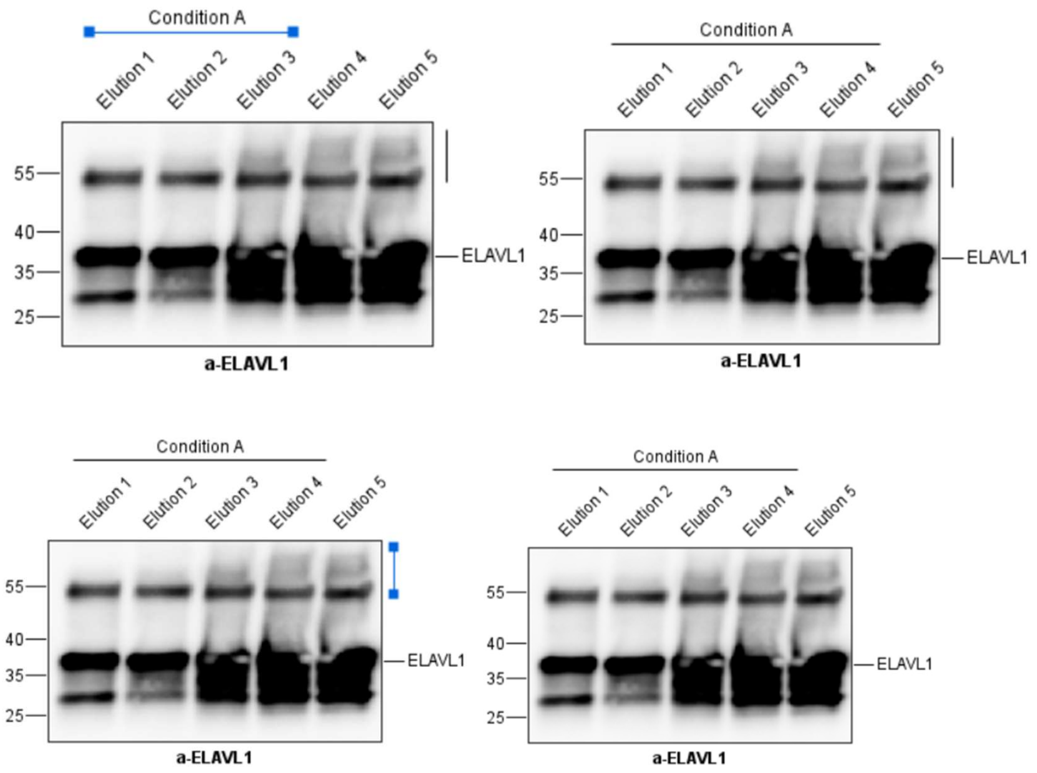

Editing band ticks, free text, and crop names

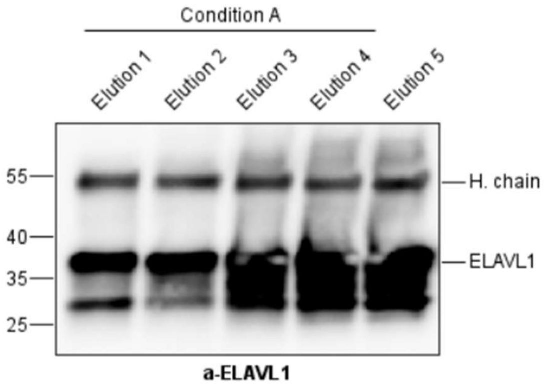

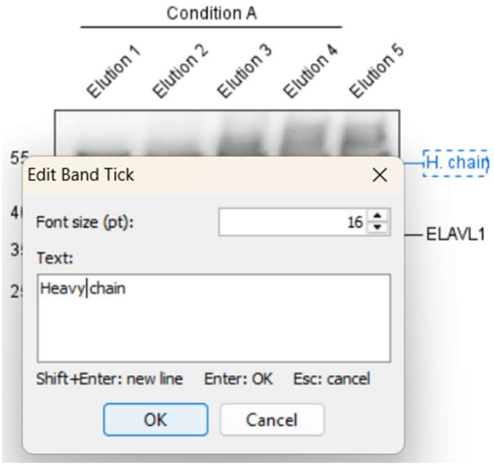

Sample labels can be edited in bulk. Multi-select the labels with shift key pressed and double click. Entered angle and font size will be applied to all selected labels.

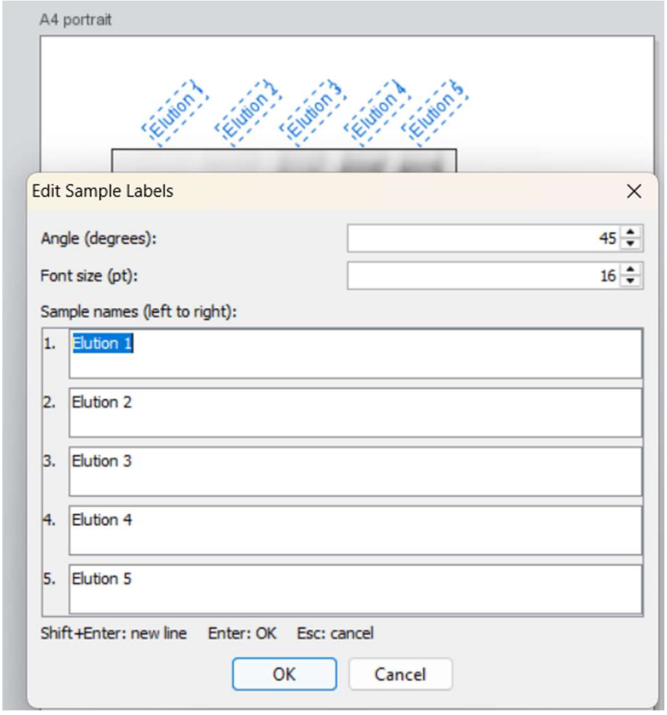

Alignment and distribution options will be shown when right-clicked.

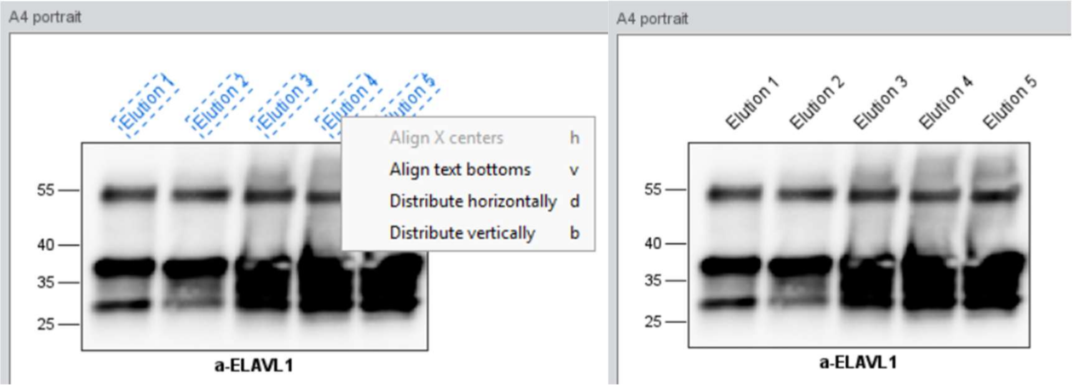

#### Saving and opening a project

At any point of figure creation, the current figure state can be saved.

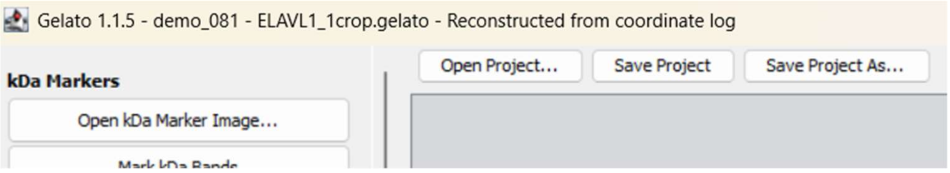

Open project… will open the figure and the source images currently used for kDa bands marking or region cropping in a new Gelato window.

#### Export

Click "Export as PDF…". All lines will be vector, all text editable. Optional check-box enables per-crop grouping of the figure elements.

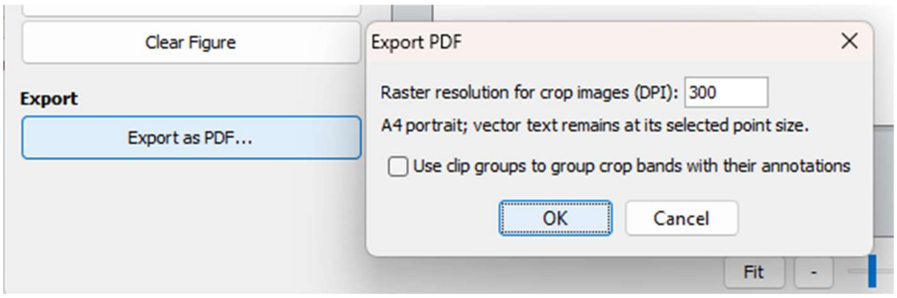

#### Save a coordinate log

1. Click Show Coordinate Log. Name the figure when prompted.
2. Click Save Log... to write a UTF-8 text file, or click Copy.

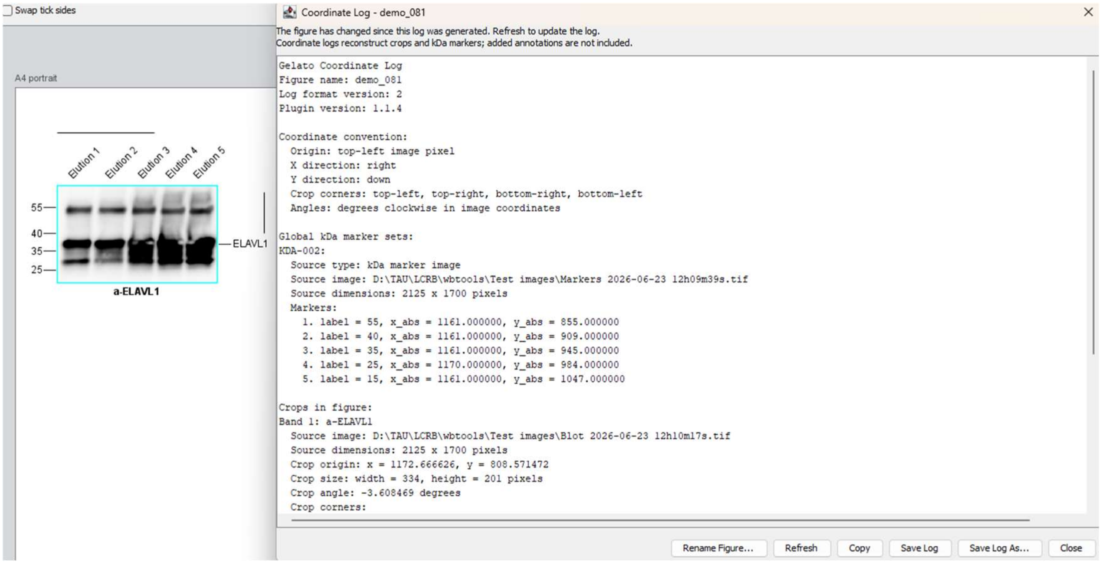

### Reconstruction and audit from a coordinate log

The coordinates log enables reproducing ROIs and Mw elements and intuitive inspection of their relationship with source images. Besides the ROI name (usually stating the protein detected) and the Mw details other text and line annotations are not recorded.

#### Reconstruct and audit

1. Click Reconstruct from Log...
2. Paste the coordinate log into the window, or click Load Log... and choose a .txt file.

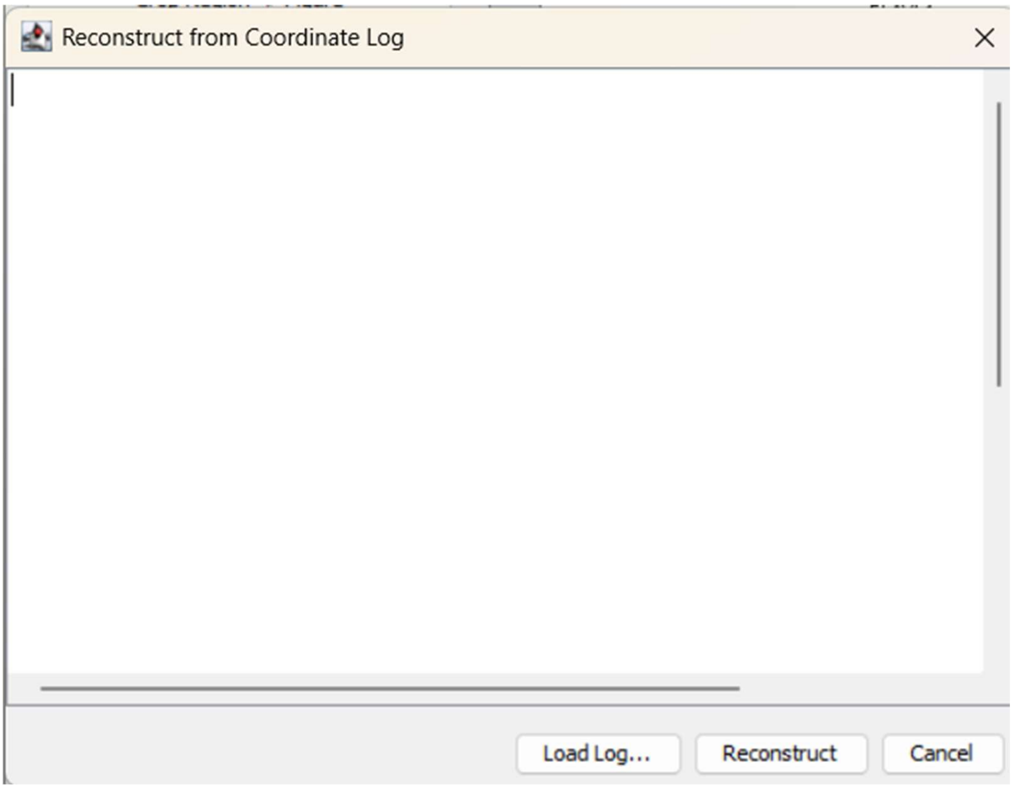

3. Click Reconstruct. If the log is incomplete or has an unsupported format version, Gelato explains.
4. Select the matching original files. The chooser shows the expected path including the file name, pixel dimensions, and how the image is used one by one. If the file exists, the chooser will pre-select it. Blot images are a required input; a separate marker-source image may be skipped.

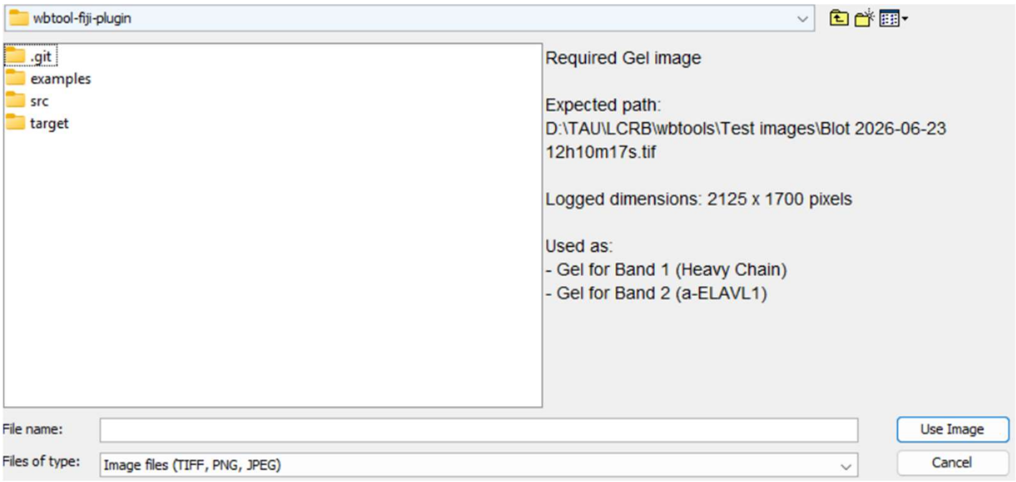

5. If the selected file has different dimensions, a warning will be shown. If the aspect ratio is unchanged, coordinates can be scaled proportionally. A different aspect ratio means X and Y are scaled separately and the result will not be trustworthy.
6. When origin/size/angle and corner coordinates disagree by more than 1 pixel, a dialogue will open. Choose which description Gelato should use, or cancel and inspect the log.
7. Inspect the reconstructed result. Gelato reconstructs all ROI panels in their logged order. It also opens annotated copies of the source images: crop outlines with band names and Mw markers appear both on gel and marker images.

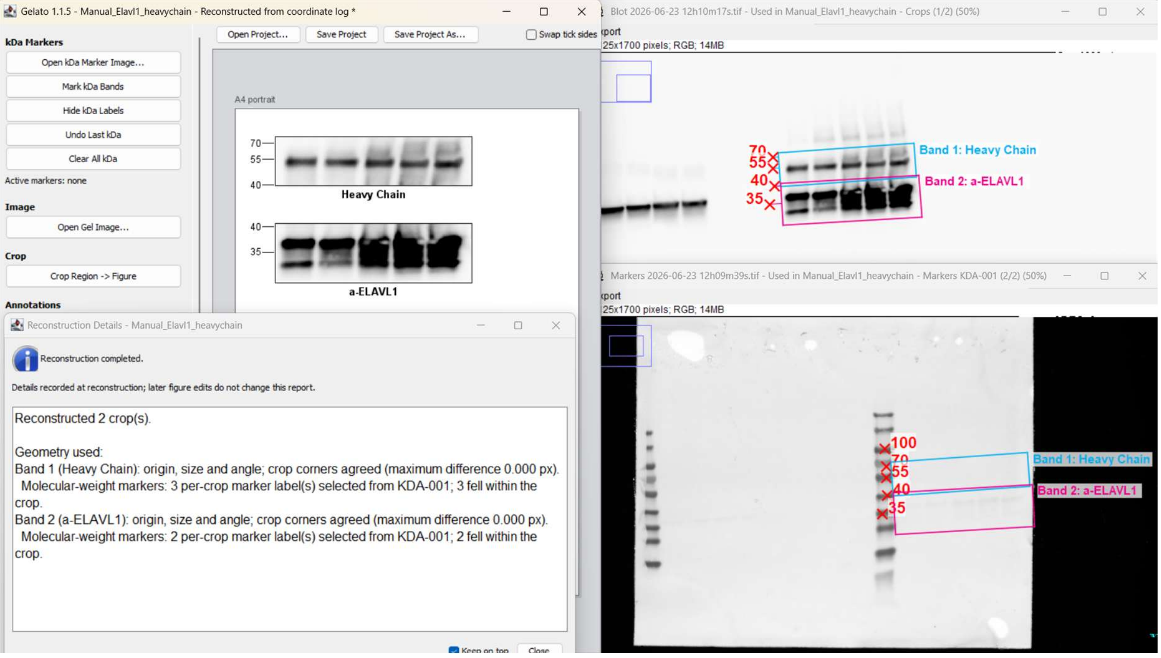

8. Reconstruction report states which crop geometry was used, how marker coordinates were resolved, and whether any source or dimension warning remains.

### How to read a coordinate log

The coordinate log is plain UTF-8 text containing the sources and the geometry needed to rebuild the figure.

#### File header

```
Gelato Coordinate Log

Figure name: Manual_Elavl1_heavychain
Log format version: 2
Plugin version: 1.1.5
```

**Gelato Coordinate Log** Identifies the file as a Gelato coordinate log.

**Figure name** can be different from the log file name. It appears on top of all windows in reconstruction mode, including source images.

**Log format version** Describes the log syntax. It is separate from the software version.

**Plugin version** Records which Gelato release created the log.

#### Coordinate convention

```
Coordinate convention:
  Origin: top-left image pixel
  X direction: right
  Y direction: down
  Crop corners: top-left, top-right, bottom-right, bottom-left
  Angles: degrees clockwise in image coordinates
```

**Origin** Absolute coordinates begin at the top-left pixel.

**X direction** X increases as you move right.

**Y direction** Y increases as you move down.

**Crop corners** Lists the fixed order used later in each crop block.

**Angles** Positive angles turn clockwise.

#### Global kDa marker sets

```
Global kDa marker sets:
KDA-001:
  Source type: kDa marker image
  Source image: D:\TAU\LCRB\wbtools\Test images\Markers 2026-06-23 12h09m39s.tif
  Source dimensions: 2125 x 1700 pixels
  Markers:
    1. label = 55, x_abs = 1161.000000, y_abs = 855.000000
```

**Global kDa marker sets** Starts the marker-set section. Only sets used by at least one crop in the figure are written.

**KDA-001** Identifier that links this marker set to one or more crops.

**Source type: kDa marker image/Gel image** States whether the markers were clicked on a separate kDa marker image or directly on a gel image

**Source image** Records the source path used when the markers were created. During reconstruction, renamed or moved file can be chosen.

**Source dimensions** Width and height of the marker image in pixels.

**Markers** Numbered list of registered markers. The label is the displayed Mw; x_abs and y_abs are the clicked source-image coordinates.

**none** Under Global kDa marker sets, this means that no marker set is used by the current figure.

#### Crops in figure

```
Crops in figure:
Band 1: a-ELAVL1
  Source image: D:\TAU\LCRB\wbtools\Test images\Blot 2026-06-23 12h10m17s.tif
  Source dimensions: 2125 x 1700 pixels
  Crop origin: x = 1172.666626, y = 808.571472
  Crop size: width = 334, height = 201 pixels
  Crop angle: -3.608469 degrees
  Crop corners:
```

```
    top-left: x = 1172.666626, y = 808.571472
    top-right: x = 1506.004450, y = 787.550167
    bottom-right: x = 1518.654996, y = 988.151672
    bottom-left: x = 1185.317172, y = 1009.172977
```

**Crops in figure** Starts the crop section.

**Band 1: a-ELAVL1** Gives the crop order and its name. Reconstruction keeps this order.

**Source image** Records the image from which this crop was taken.

**Source dimensions** Records the full source-image dimensions.

**Crop origin** Gives the top-left corner of the rotated crop in source-image coordinates.

**Crop size** Gives the crop width and height before it is scaled for display on the A4 canvas.

**Crop angle** Gives the crop rotation in degrees using the convention above.

**Crop corners** Gives the four absolute corner coordinates. This can be used for reconstruction instead of origin, size and angle, because it's easier to provide corner coordinates when the figure is constructed in a different program.

**none** If logs contains no crops, the figure cannot be reconstructed.

#### Used kDa markers

```
  Used kDa markers:
    Marker set: KDA-001
    Marker source image: D:\TAU\LCRB\wbtools\Test images\Markers 2026-06-23 
12h09m39s.tif
    Marker source dimensions: 2125 x 1700 pixels
    Gel dimensions: 2125 x 1700 pixels
    Coordinate scale: x = 1.000000, y = 1.000000
    1. label = 55, source_x_abs = 1161.000000, source_y_abs = 855.000000, gel_x_abs = 
1161.000000, gel_y_abs = 855.000000, y_in_crop = 45.602206
```

**Marker set** Names the global set used for this crop. Can be none.

**Marker source image** Repeats the marker source of the marker set for convenience.

**Marker source dimensions** Repeats the marker-image dimensions used for this mapping.

**Gel dimensions** Dimensions of the crop source image.

**Coordinate scale** Shows the X and Y multipliers used to transfer marker coordinates into the gel image. A value of 1 means no scaling in that direction.

**label** Identifies the marker selected from the global set.

**source_x_abs / source_y_abs** Original click position on the marker source.

**gel_x_abs / gel_y_abs** The click position after scaling into the gel coordinate system.

**y_in_crop** Marker height measured downward from the crop top edge in the crop's rotated coordinate system. This is the value used to place the tick.

**Detailed marker coordinates** Serve as an audit trail. During reconstruction, Gelato recalculates them from the global marker set. If a supplied value differs by more than 1 pixel, Gelato warns and uses the recalculated value.

**WARNING or Note** Appears when marker and gel dimensions differ. A warning means their aspect ratios differ; a note means the aspect ratio is preserved.

**No markers from this set fell within the crop** Means the set is linked, but none of its marker heights lies inside this crop.

#### If you edit or create a log by hand

1. Keep the exact section names, field names, separators, and decimal points shown in a Gelato-generated log.
2. For every Marker set:
    a. Give a source image name or path. If you include source dimensions, it can be checked later.
    b. Enter markers values and positions.
3. For every crop:
    a. Give a source image name or path. If you include source dimensions, it can be checked later.
    b. Describe crop geometry either with Crop origin (top left corner) and Crop size (Crop angle may be omitted and then defaults to 0 degrees), or with at least three labeled crop corners.
    c. Molecular-weight information is optional. If a crop names a marker set but has no per-crop marker list, Gelato evaluates every marker in that set and keeps those within the crop.

### How coordinates are handled

Gelato keeps the original click coordinates and derives each later coordinate from them. This makes the mapping reproducible and lets the reconstruction recalculate, rather than simply trust, detailed entries in the log.

#### 1. Marker source to gel

A marker is first stored at an absolute source position (x_source, y_source). For a separate marker image, Gelato compares the marker-image dimensions with the gel dimensions:

```
scale_x = gel_width / marker_width
scale_y = gel_height / marker_height
x_gel = x_source * scale_x
y_gel = y_source * scale_y
```

When both images have the same dimensions, both scale values are 1. If the dimensions differ but the aspect ratio matches, the transfer is proportional. If the aspect ratios differ, X and Y are scaled independently and Gelato records a warning.

#### 2. Gel coordinates to the rotated crop

Let (x0, y0) be the crop origin and theta its clockwise angle. Gelato subtracts the origin, then rotates the marker into the crop's local coordinate system.

```
dx = x_gel - x0
dy = y_gel - y0
x_in_crop = cos(theta) * dx + sin(theta) * dy
y_in_crop = -sin(theta) * dx + cos(theta) * dy
```

Gelato needs y_in_crop for molecular-weight placement, and that value is written to the log. A tilted crop therefore keeps each tick aligned with the corresponding source height.

#### 3. Crop inclusion and tick placement

A marker is assigned to a crop when its local vertical position lies between -0.5 pixel and crop_height + 0.5 pixel.

```
include marker when -0.5 <= y_in_crop <= crop_height + 0.5
tick_y_on_canvas = crop_top_on_canvas + round(y_in_crop * crop_display_scale)
```

When you make a crop narrower or wider on the canvas, crop_display_scale changes. The tick moves with the crop, while the logged source position remains unchanged.

#### 4. Crop corners

The origin, size, and angle also determine the four logged corners:

```
top_left     = (x0, y0)
top_right    = (x0 + width*cos(theta), y0 + width*sin(theta))
bottom_left  = (x0 - height*sin(theta), y0 + height*cos(theta))
bottom_right = top_right + bottom_left - top_left
```

During reconstruction, Gelato can rebuild a crop from origin/size/angle or from at least three labeled corners. If all descriptions are present and disagree by more than 1 pixel, it asks the user which geometry to use.

#### 5. Scaling during reconstruction

If you choose an image whose pixel dimensions differ from those recorded in the log, Gelato first calculates selected_scale_x = selected_width / logged_width and selected_scale_y = selected_height / logged_height. It scales the crop origin and the neighboring corner directions, then recomputes the width, height, and angle. With a matching aspect ratio this is proportional. With a different aspect ratio, the geometry may change, so the result is flagged as potentially inexact.

#### Example from the demo log

The demo marker and gel images are both 2125 x 1700 pixels, so scale_x = scale_y = 1. Marker 40 stays at (1164, 900). For Band 1, the crop origin is (1178.812988, 892.642273) and the angle is -3.483271 degrees. Substituting those values gives y_in_crop = 6.444140 pixels, matching the log.

### Example log

```
Gelato Coordinate Log
Figure name: Manual_Elavl1_heavychain
Log format version: 2
Plugin version: 1.1.5
Coordinate convention:
  Origin: top-left image pixel
  X direction: right
  Y direction: down
  Crop corners: top-left, top-right, bottom-right, bottom-left
  Angles: degrees clockwise in image coordinates

Global kDa marker sets:
KDA-001:
  Source type: kDa marker image
  Source image: D:\TAU\LCRB\wbtools\Test images\Markers 2026-06-23 12h09m39s.tif
  Source dimensions: 2125 x 1700 pixels
  Markers:
    1. label = 55, x_abs = 1161.000000, y_abs = 855.000000
    2. label = 40, x_abs = 1161.000000, y_abs = 909.000000
    3. label = 35, x_abs = 1161.000000, y_abs = 945.000000
    4. label = 25, x_abs = 1170.000000, y_abs = 984.000000
    5. label = 15, x_abs = 1161.000000, y_abs = 1047.000000

Crops in figure:
Band 1: a-ELAVL1
  Source image: D:\TAU\LCRB\wbtools\Test images\Blot 2026-06-23 12h10m17s.tif
  Source dimensions: 2125 x 1700 pixels
  Crop origin: x = 1172.666626, y = 808.571472
  Crop size: width = 334, height = 201 pixels
  Crop angle: -3.608469 degrees
  Crop corners:
    top-left: x = 1172.666626, y = 808.571472
```

```
    top-right: x = 1506.004450, y = 787.550167
    bottom-right: x = 1518.654996, y = 988.151672
    bottom-left: x = 1185.317172, y = 1009.172977
  Used kDa markers:
    Marker set: KDA-001
    Marker source image: D:\TAU\LCRB\wbtools\Test images\Markers 2026-06-23
12h09m39s.tif
    Marker source dimensions: 2125 x 1700 pixels
    Gel dimensions: 2125 x 1700 pixels
    Coordinate scale: x = 1.000000, y = 1.000000
    1. label = 55, source_x_abs = 1161.000000, source_y_abs = 855.000000, gel_x_abs =
1161.000000, gel_y_abs = 855.000000, y_in_crop = 45.602206
    2. label = 40, source_x_abs = 1161.000000, source_y_abs = 909.000000, gel_x_abs =
1161.000000, gel_y_abs = 909.000000, y_in_crop = 99.495147
    3. label = 35, source_x_abs = 1161.000000, source_y_abs = 945.000000, gel_x_abs =
1161.000000, gel_y_abs = 945.000000, y_in_crop = 135.423775
    4. label = 25, source_x_abs = 1170.000000, source_y_abs = 984.000000, gel_x_abs =
1170.000000, gel_y_abs = 984.000000, y_in_crop = 174.912898
```
