## Supplementary figures and images for "Gelato streamlines reproducible and auditable assembly of western blot figures"

### Supplemental Figure 1

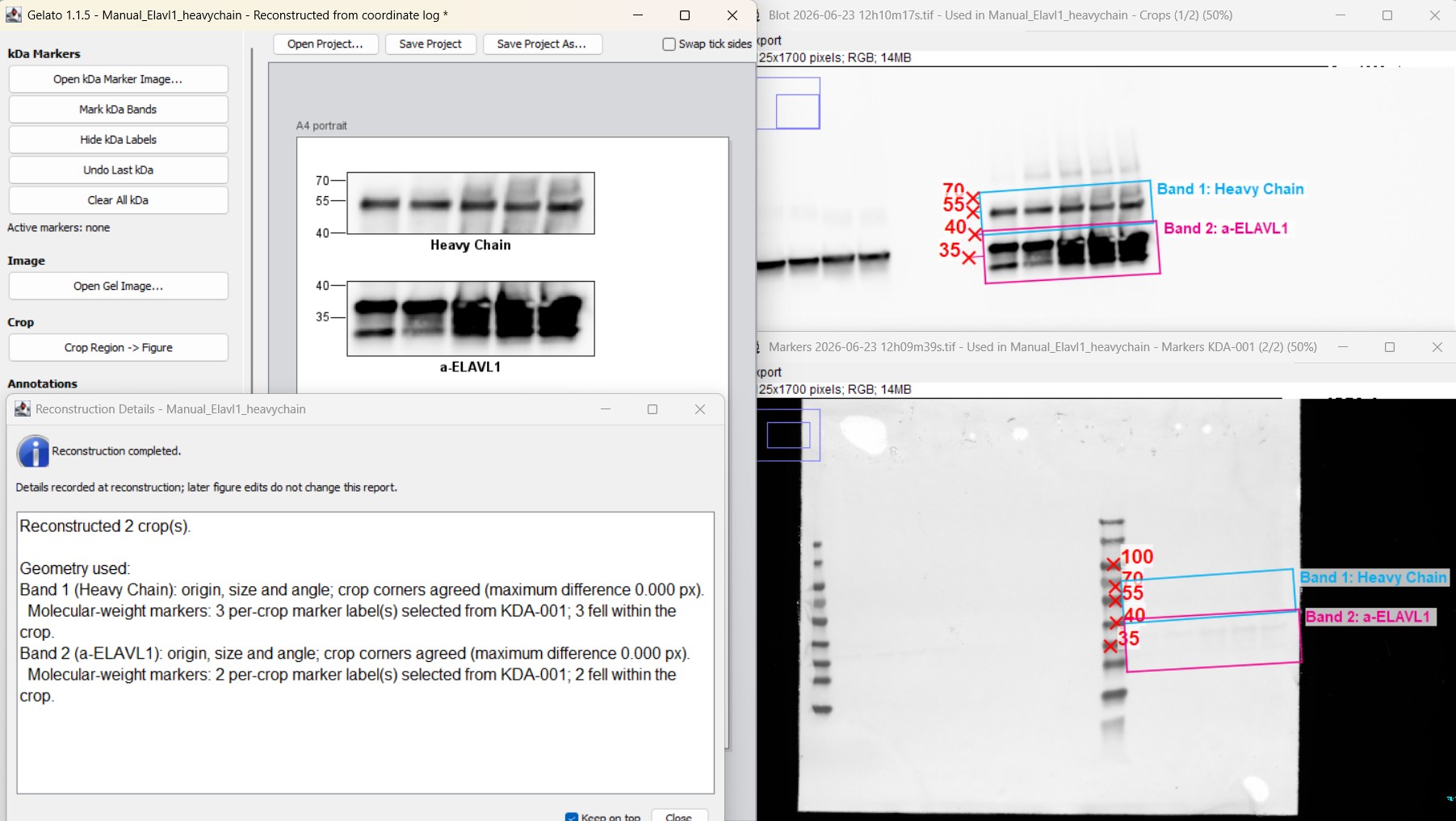
