## Supplemental Table 1 for "Gelato streamlines reproducible and auditable assembly of western blot figures"

|  | Gelato | QuickFigures | FigureJ | LabFigures (Sciugo) | Bio-Rad Image Lab | Adobe Illustrator |
| --- | --- | --- | --- | --- | --- | --- |
| Primary scope | Gel figure assembly | Figure assembly | Multi-panel figure assembly | Figure assembly | Gel/blot acquisition and analysis | Figure assembly |
| Approach for Integrity and Reproducibility | Reconstruction from human-readable coordinate and crop report | File paths export | Access to raw data and processing history | - | Metadata | - |
| Molecular weight annotations | Mapped from colorimetric image on blot image | Manually positioned after ROI is cropped | Manual | Marked on combined image before ROI is cropped | Automatic MW bands recognition on combined image | Manual |
| Independent ROI assembly | Rotation per crop | Rotation per crop | Rotation per figure | Rotation per crop | Not intended | Manual |
| Annotations on exported pdf figure | Fully editable vector | Fully editable vector | Fully editable vector | Vector | Not supported | Fully editable vector |

*Supplementary Table 1.* Comparison of Gelato with existing software used for western blot figure preparation. Tools are compared by primary scope, features supporting integrity and reproducibility, molecular-weight annotation workflow, assembly of multiple ROIs to one figure, and flexibility of annotations on exported figures.
